# Gut distension evokes rapid neural dynamics in vagal and hindbrain populations of larval zebrafish

**DOI:** 10.1101/2025.07.21.665834

**Authors:** Minel Arinel, John A. Atkinson, Karina M. Matos-Fernández, Evan P. Drage, Matt Hawkyard, Eva A. Naumann

## Abstract

Animals sense food quantity and quality to regulate feeding behaviors essential for survival. Enteroendocrine cells (EECs) in the gut epithelium detect luminal distension and nutrients, signaling this information to the brain via vagal sensory neurons. However, how mechanosensory and chemosensory signals are dynamically encoded by gut-brain circuits remains unclear, particularly during early development. Leveraging the transparency and genetic tractability of larval zebrafish, we developed a feeding assay offering liposome-based complex particles designed to release specific nutrients after consumption. By quantifying gut fluorescence after exposure to these particles in 9-day-old zebrafish, we demonstrate that EECs regulate nutrient-specific feeding soon after functional gut formation. To determine how these post-ingestive signals are encoded along the gut-brain circuitry, we developed a microgavage method enabling simultaneous stimulus delivery to the gut and volumetric two-photon calcium imaging. We found that repeated gut distension alone drove widespread activation and suppression in vagal and dorsal hindbrain neurons, emerging as a dominant visceral signal after two days of feeding onset. Although nutrient-evoked responses shared temporal features with distension, allyl isothiocyanate (AITC), an aversive compound found in wasabi, elicited neural dynamics with slower onset kinetics. These findings reveal that fast gut-to-brain communication emerges early in life, with mechanical distension and aversive chemical cues encoded through distinct temporal dynamics in developing interoceptive circuits.

**Highlights:**

- Larval zebrafish enteroendocrine cells regulate nutrient-specific feeding
- Left and right vagus similarly encode gut distension and nutrients
- Gut distension drives rapid, widespread neural responses across the hindbrain
- Enteric stimuli evoke diverse fast neural dynamics in early development

## Introduction

Animals must rapidly evaluate both the quantity and quality of ingested food to make adaptive feeding decisions essential for survival. While pre-ingestive cues like taste and smell initiate feeding, the gut provides definitive confirmation of how much and what has been consumed^1,2^. This post-ingestive information is transmitted within seconds to coordinate digestion, prevent toxin ingestion, and regulate satiety^3–6^. Although recent research has highlighted the importance of gut-brain communication in adult mammals^6–15^ and insects^16–27^, how these circuits develop and function in early life remains poorly understood^28,29^.

Central to post-ingestive signaling are enteroendocrine cells (EECs), specialized sensors comprising ∼1% of intestinal epithelial cells that detect luminal distension and nutrients^30–32^. EECs promote food intake during hunger, reduce intake after nutrient detection, and trigger aversive responses to harmful compounds^33–35^. A specialized subset of EECs, termed neuropods, forms synaptic connections with vagal sensory neurons that innervate the gut^1,2,12,32,33,36–41^. These vagal afferents rapidly transmit mechanosensory and chemosensory information to the brain via both EEC signaling and their intrinsic receptors^12,14,15,40,42,43^. Vagal afferents then project to the nucleus tractus solitarius (NTS) and area postrema (AP) in the brainstem^6,44–46^, which further relay this information to higher-order brain regions^6,40,47^ and to the dorsal motor nucleus of the vagus^6,48,49^, completing a bidirectional feedback loop to modulate peripheral physiology.

In adult mammals, vagal sensory neurons integrate mechanical and nutrient cues in a concentration-dependent manner^50^ and exhibit heterogeneous activity kinetics^51–53^. They also show molecular^54^, anatomical^6,55^, and functional^6,55^ lateralization between the left and right vagal branches. In the brainstem, NTS neurons can be activated or suppressed depending on the stimulated gastrointestinal organ, displaying diverse response durations to gut distension^10^. However, most studies rely on anesthetized preparations, limiting understanding of dynamic gut-brain signaling in awake, developing organisms^56^. This gap is particularly significant because while the anatomical formation of these circuits occurs as early as embryonic day 16.5 in mice^57^, the temporal encoding strategies that underlie this rapid gut-brain communication and how they emerge functionally during early postnatal development remain largely unexplored^58^.

Larval zebrafish offer unprecedented experimental access to investigate developing vertebrate gut-brain circuits. Unlike mammals, zebrafish develop externally, allowing direct observation and manipulation at early ages^59^. By 5 days post-fertilization (dpf), they have resorbed their yolk, developed a functional gut containing EECs, and initiated exogenous feeding^60,61^. Importantly, zebrafish share key features of the mammalian gut-brain axis, including molecularly defined EEC subtypes, vagal innervation of the gut, and homologous brainstem circuits^60–66^. Their optical transparency and small size enable access to both the gut and the brain, making them ideal for dissecting gut-brain signaling *in vivo* with single-cell resolution.

Previous studies in larval zebrafish have shown that EECs and vagal neurons express receptors sensing mechanical (Piezo2) and chemical (including Sglt1, Glyt1, Trpa1) stimuli and exhibit stimulus-evoked responses^62,63,67–69^. Zebrafish EECs form basal projections resembling mammalian neuropods^62,63^. Additionally, zebrafish vagal neurons are activated by intraluminal delivery of glucose^70^, the Trpa1 agonist allyl isothiocyanate^63^ (AITC; a pungent compound found in wasabi), or other microbial byproducts, suggesting that gut-brain circuits are functionally active at larval stages. However, whether these early anatomical connections translate into feeding-relevant signaling and how enteric information is rapidly encoded in the developing nervous system are poorly understood.

Here, we developed two complementary approaches to investigate gut-brain signaling in larval zebrafish. First, we exposed hungry larval zebrafish to alginate-encapsulated, liposome-based complex particles (CPs)^71^ that release nutrients and fluorescein after ingestion, thereby isolating gut-derived responses from exteroceptive cues. By quantifying gut fluorescence after offering these food-sized CPs, we show that amino acids promoted consumption while glucose had no effect. Chemogenetic EEC depletion revealed their dual effect, reducing food approach probability while paradoxically increasing nutrient-specific consumption once feeding began, implicating gut-brain signaling in early feeding decisions.

Second, to investigate how these gut-derived signals are represented in the brain, we developed a prism-guided microgavage method compatible with two-photon calcium imaging in awake zebrafish larvae. This approach enabled precise enteric stimulation while simultaneously recording neural activity in real time. We mapped stimulus-evoked activity across the bilateral vagal sensory ganglia and dorsal hindbrain, including the vagal sensory lobe, homologous to the mammalian NTS^72^, and vagus motor nucleus, homologous to the dorsal motor nucleus of the vagus^73^.

These findings establish that zebrafish vagal neurons encode fast, dynamic information about gut distension with minimal lateralization, distributing this enteric information across the hindbrain as soon as the gut-brain circuitry becomes functional. Rather than developing gradually, the capacity for rapid visceral signaling appears to be a feature of the earliest feeding behaviors. Compared to nutrient-evoked activity, distension emerged as a dominant driver of vagal and hindbrain responses in early development. Together, our results show that larval zebrafish is a tractable model for investigating rapid gut-brain communication and demonstrate that distension and aversion are dynamically encoded in developing interoceptive circuits.

## Results

### Amino acids increase gut-mediated feeding

To determine how nutrient signaling from the gut regulates feeding, we encapsulated specific nutrients within complex particles^71,74^ (CPs), designed to bypass exteroceptive chemosensing and restrict exposure to the gastrointestinal tract (**Figure 1A**). To form CPs (**Figure 1B**), target nutrients were combined with sodium fluorescein in an aqueous solution, encapsulated within liposomes, and then embedded in an alginate matrix, which is considered tasteless and odorless^75^. As CPs show high retention of water-soluble compounds when suspended in water^71^, they expose their encapsulated nutrients to the gut epithelium upon ingestion, where EECs may detect and relay nutrient-specific signals to the brain.

**Figure 1.**
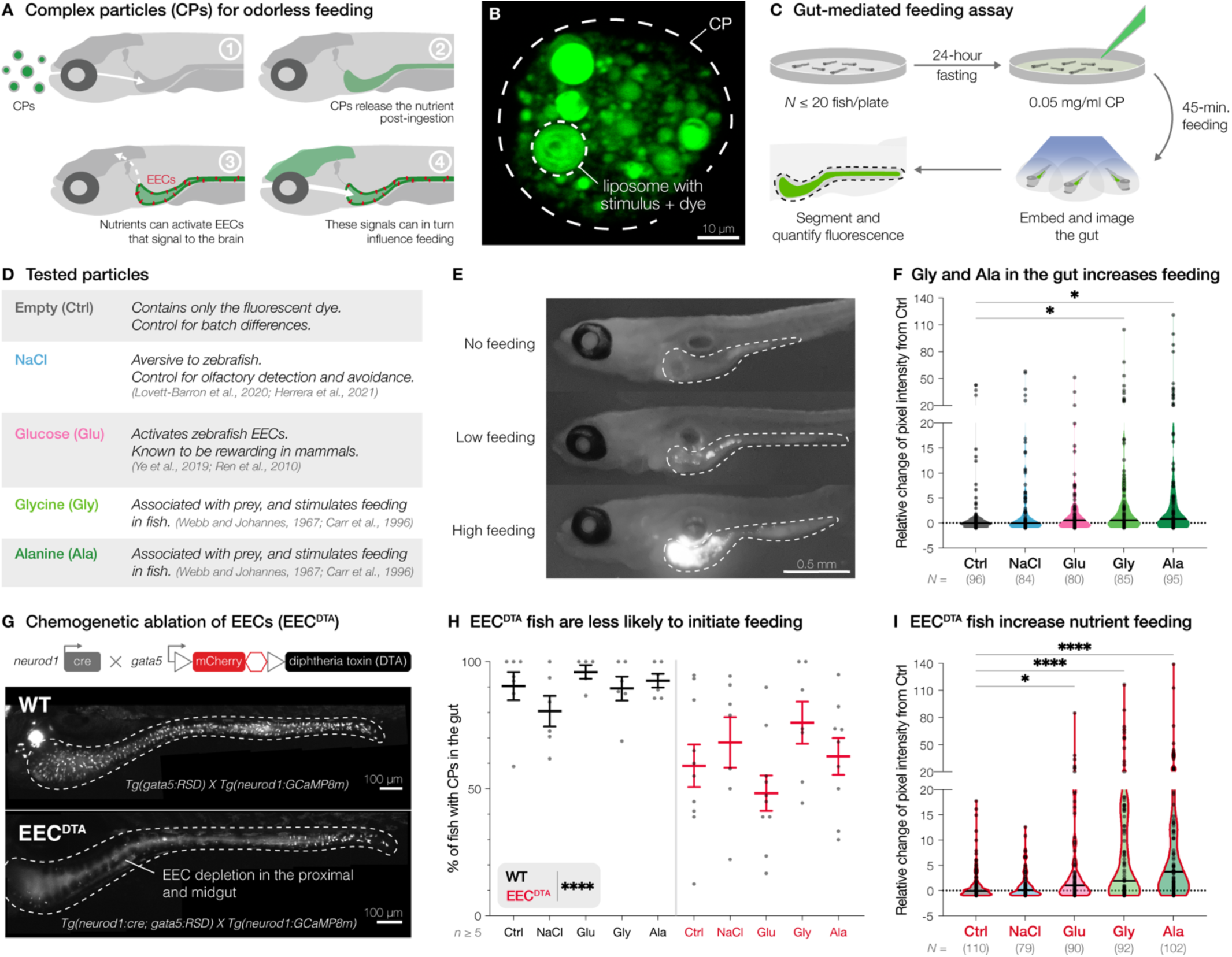
Zebrafish EECs mediate nutrient-specific feeding. (**A**) Schematic illustrating gut nutrient delivery by feeding complex particles (CPs), which bypass exteroceptive chemosensation as they expose their contents only in the gut, potentially activating EECs (red). (**B**) Two-photon image of a CP with fluorescent liposomes surrounded by alginate matrix (dashed outline). (**C**) After 24 hours of fasting, freely swimming zebrafish were exposed to CPs loaded with different nutrients. After 45 minutes of feeding, zebrafish were fixed to quantify gut fluorescence. (**D**) Table of tested nutrients and rationale for selection. (**E**) Side views of representative zebrafish with no, low, and high consumption of fluorescent CPs in the gut (dashed outlines). (**F**) Gut fluorescence in wild-type (WT) zebrafish, plotted as relative change from the median fluorescence of the batch control. Compared to empty CPs (Ctrl, *N* = 96) CPs with glycine (Gly, *N* = 85, \**P* = 0.0155) and alanine (Ala, *N* = 95, \**P* = 0.0119) drove significant increase in feeding, unlike NaCl (*N* = 84, *P* > 0.9999) or glucose (Glu, *N* = 80, *P* = 0.7261, Kruskal-Wallis and Dunn’s multiple comparisons tests). Dots represent individual fish. (**G**) Top: schematic of chemogenetic EEC ablation. Bottom: representative gut images of Cre-negative and Cre-positive EEC-depleted fish (EEC^DTA^). (**H**) Percentage of fish with any gut fluorescence in each batch of WT and EEC^DTA^ zebrafish across stimuli (*n* ≥ 5 plates). EEC^DTA^ exhibited significantly lower % of fish with CPs (\*\*\*\**P* < 0.0001, Kolmogorov-Smirnov test) with higher variability (\*\*\*\**P* < 0.0001, Fligner-Killeen test) than WT. Dots represent experimental batches. (**I**) Gut fluorescence in EEC^DTA^ zebrafish fed with CPs, plotted as relative change from the median fluorescence of the batch control. Compared to empty CPs (*N* = 110), EEC^DTA^ zebrafish ate significantly more CPs filled with glucose (*N* = 90, \**P* = 0.0163), glycine (*N* = 92, \*\*\*\**P* < 0.0001), and alanine (*N* = 102, \*\*\*\**P* < 0.0001), but not with NaCl (*N* = 79, *P* > 0.9999, Kruskal-Wallis and Dunn’s multiple comparisons tests). Dots represent individual fish. Violin plots show median. Scatter plot shows mean ± SEM.

Zebrafish were fed with live paramecia from 4–7 dpf and then fasted for 24 hours (**Methods**). At 9 dpf, sibling batches were exposed to CPs (0.05 mg/L, 25 mM nutrient concentration; **Figure 1C**). After 45 minutes of free feeding, these zebrafish were euthanized, fixed, and individually embedded on their side to quantify particle consumption by measuring gut fluorescence.

To test how compounds released in the gut affect feeding, we designed different CPs (**Figure 1D**), including empty particles containing only sodium fluorescein as a negative control (Ctrl; **Figure S1**). Sodium chloride (NaCl) particles acted as an additional negative control from CP nutrient leaching, since zebrafish avoid NaCl via olfactory pathways^76,77^. For nutrients, we tested glucose (Glu), since it activates zebrafish EECs^62,63^ and is highly rewarding in mammals^78–80^. Additionally, we generated CPs containing the amino acids glycine (Gly) and alanine (Ala) as they increase feeding across various fish species and are abundant in their natural prey^81,82^.

To quantify the consumption of different CPs, we measured total gut fluorescence in individual zebrafish (**Figures 1E** and **S1A**). Due to significant batch-to-batch variation with Ctrl particles (**Figure S1B**), we normalized each fish’s fluorescence value to its respective median batch Ctrl intensity. As expected, NaCl particles did not change feeding relative to Ctrl, confirming that consumption is not affected by nutrient leaching. Compared to the Ctrl group, zebrafish exposed to glycine or alanine, but not glucose, significantly increased consumption (**Figure 1F**). These results demonstrate that exposure of glycine and alanine to the gut promotes feeding, underscoring the ecological importance of amino acids in the zebrafish diet.

### EEC depletion alters food approach and nutrient-specific feeding

To test whether gut-mediated feeding behaviors are modulated by EECs, we chemogenetically depleted EECs from the proximal and midgut using diphtheria toxin^63^ (EEC^DTA^; **Figure 1G**) and exposed EEC^DTA^ zebrafish to the same CPs as wild-type (WT) cohorts. Examining whether EEC depletion altered the likelihood of food approach, we quantified the percentage of zebrafish containing any fluorescent particles in their gut. EEC^DTA^ batches had significantly lower and more variable food approach compared to WT across stimuli (**Figure 1H**). However, in EEC^DTA^ zebrafish, glucose, glycine, and alanine each significantly increased consumption compared to Ctrl, whereas NaCl did not (**Figure 1I**). Notably, the relative intensities of nutrients were greater in EEC^DTA^ zebrafish compared to WT (glucose: ∼1.8x, glycine: ∼3.2x, alanine: ∼4.7x). To determine whether this increased fluorescence in EEC^DTA^ fish was caused by impaired peristalsis, we compared gut motility between unfed and fed EEC^DTA^ zebrafish relative to WT and Cre-negative controls (**Figure S2** and **Video S1**). As expected, fed zebrafish exhibited significantly higher gut motility compared to unfed fish, but no differences in gut motility were detected across genotypes. These results, together with the nutrient-specific increase in fluorescence in EEC^DTA^ fish, suggest that the elevated fluorescence reflects greater nutrient intake rather than impaired gut clearance. Collectively, these findings reveal a dual role for EECs in modulating feeding behavior. While EEC depletion reduced the likelihood of initiating feeding, it enhanced nutrient intake once feeding began. These results suggest that even in early development, zebrafish EECs regulate nutrient-specific feeding by conveying satiety-related signals to the brain.

### Microgavage during neural recording of gut-brain signaling

To investigate how these enteric signals are encoded by the nervous system, we next aimed to record gut-derived neural responses, focusing on signals transmitted via the vagus nerve to the brain. Intriguingly, larval zebrafish EECs exhibit basal protrusions morphologically resembling neuropods that form synapses in mammals^63^ for fast information transmission (**Figure 2A**). However, not much is known about rapid gut-brain communication at early developmental stages^57^, as recordings are challenging in mammals during development^83,84^.

**Figure 2.**
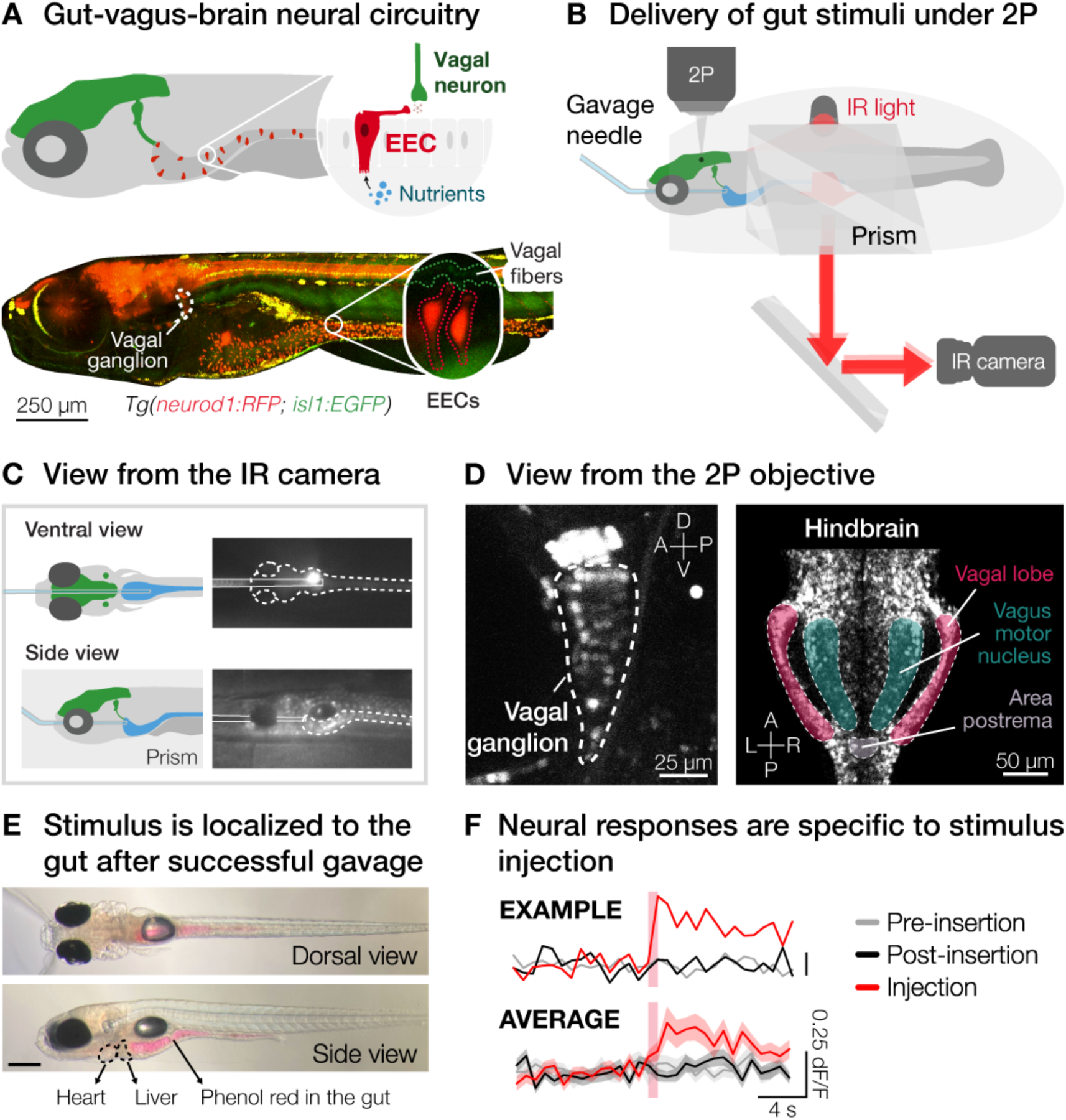
Microgavage during two-photon imaging to record gut-mediated neural activity. (**A**) Top: schematic of the gut-vagus-brain neural circuitry. Bottom: two-photon image of a 7 dpf *Tg(neurod1:RFP; isl1:EGFP)* zebrafish, showing the entire EEC population (red) and vagal neurons (green), with a zoom-in of two EECs with protrusions toward vagal fibers. Schematic of the microgavage setup with zebrafish embedded in agarose for imaging. A bent gavage needle is guided into the gut using a prism mirror for simultaneous ventral and side visualization, ensuring precision enteric stimulus delivery. (**B**) Left: cartoons of the infrared (IR) camera’s field of view. Right: snapshots of ventral and side views for precise microgavage needle positioning. (**C**) Left: two-photon image of a left vagal ganglion (dashed outline). The entire vagal ganglion can be imaged within the field-of-view. Scale bar = 25 µm. Right: two-photon image of a plane in the dorsal hindbrain. The vagal sensory lobe, area postrema (located more dorsally than the shown plane), and vagus motor nucleus can be imaged simultaneously in our imaging volume. A, anterior; P, posterior; D, dorsal; V, ventral; L, left; R, right. (**D**) Transmission images after successful microgavage show enteric stimulus localization exclusively in the gut (red), confirming intact internal organ morphology (e.g., heart and liver). Scale bar: 250 µm. (**E**) Neurons show time-locked responses specifically to enteric stimulation, not to the microgavage procedure. dF/F traces of a representative vagal neuron (top) and average population response (bottom). Red vertical bars indicate the time of injection. Mean ± SEM.

Adapting a previous method for microgavaging anesthetized larval zebrafish^85^, we developed a novel microgavage technique to deliver nanoliters of stimuli into the gut during simultaneous two-photon calcium imaging in awake larval zebrafish (**Figures 2B** and **S3**). To eliminate the need for anesthesia, we designed a flexible microgavage needle from aluminosilicate glass with an extended taper and a fine tip (≤20 µm outer diameter; **Figure S3C**), allowing gentle insertion through the pharynx and esophagus without breakage. A ∼30° bend near the tip enabled precise alignment with the gastrointestinal tract, ensuring accurate stimulus delivery and an optimal imaging angle. To permit simultaneous visualization from ventral and lateral perspectives, we placed a prism next to the 6–10 dpf zebrafish embedded in 1.5% low-melting-point agarose (**Figures 2C**, **S3A**, and **S3B**).

To perform two-photon calcium imaging during microgavage, we used *Tg(elavl3:H2B-GCaMP8m)* zebrafish, which express the nuclear-localized calcium indicator GCaMP8m, in two imaging configurations, optimized to record neural activity in either vagal ganglia or dorsal hindbrain (**Figures 2D, S3A**, and **S3B**). In each configuration, the microgavage needle is guided through the gastrointestinal tract while monitoring for any signs of discomfort or pain, such as twitching, via an infrared camera (**Video S2**). After each experiment, we visually confirmed enteric stimulus localization in the gut and verified that the procedure did not cause injury (**Figure 2E**).

To validate that neural responses were driven by gut stimulation rather than the microgavage procedure, we compared neural activity prior to needle insertion, post-insertion without stimulus injection, and after intraduodenal stimulus delivery (**Figure 2F**). Confirming minimal abrasion, any observed neuronal activity was stimulus-dependent and time-locked to gut stimulus injection. Therefore, our microgavage method provides the necessary temporal and spatial precision to reliably capture fast gut-derived neural signals, confirming functional gut-to-brain connectivity in larval zebrafish as early as 6 dpf.

### Vagal afferents bilaterally encode gut distension

Since each microgavage injection inherently induces gut distension, we first investigated vagal responses to stretch stimuli without the presence of nutrients. Recording neural activity in either the left or right vagus, we injected the standard E3 medium in which zebrafish are raised (**Methods**), hereafter referred to as “water”, directly into the gut lumen in five pulses of 2 nL, each separated by a 2-minute interval (**Figure 3A**).

**Figure 3.**
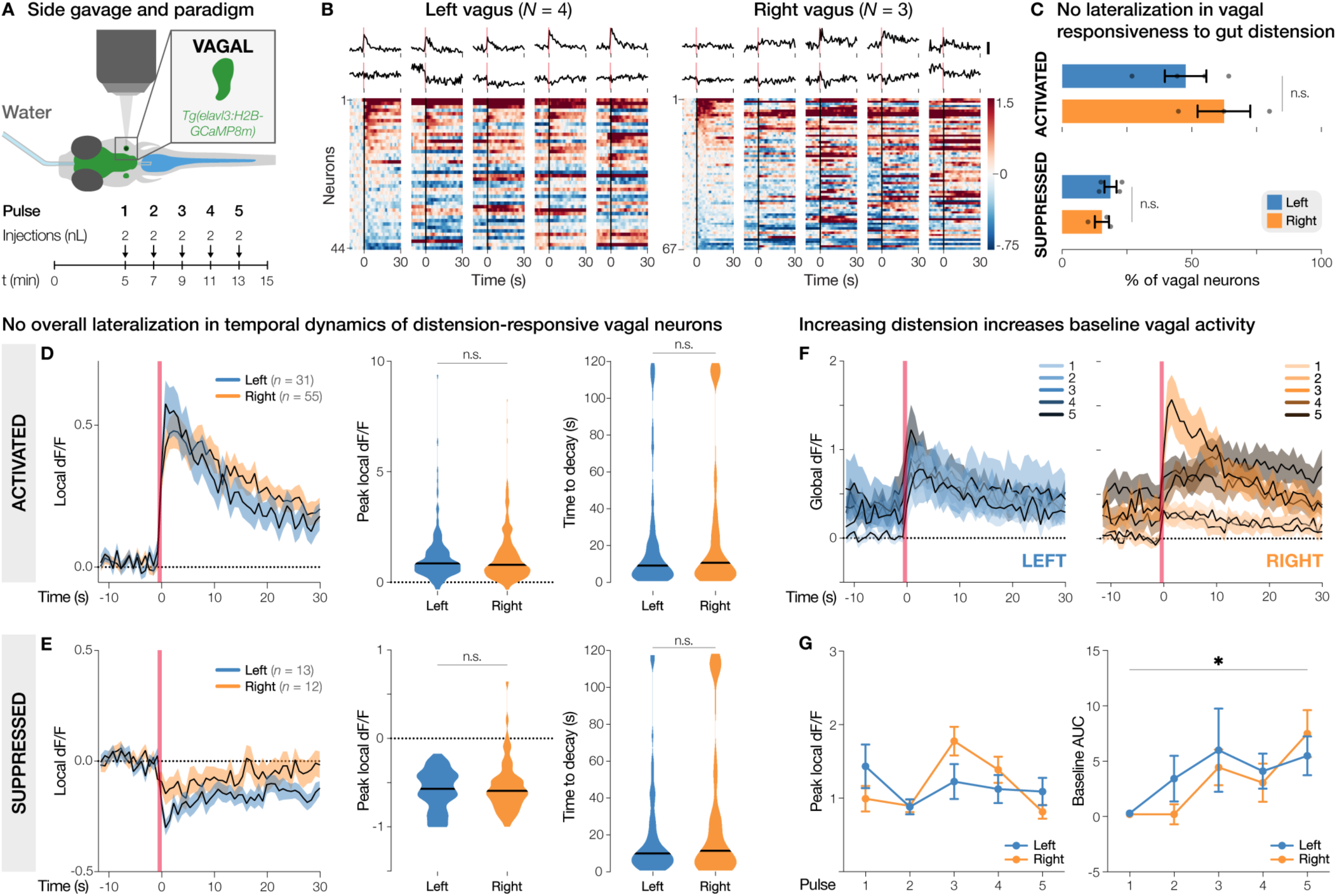
Gut distension elicits bilateral activation and suppression of vagal neurons. (**A**) Experimental setup. 6–10 dpf *Tg(elavl3:H2B-GCaMP8m)* zebrafish were embedded on their side and continuously imaged during microgavage with E3 medium (“water”), delivering five pulses of 2 nL. (**B**) Top: Normalized dF/F traces across five pulses from representative activated (top) and suppressed (bottom) neurons in the left and right vagal ganglia. Scale bar: 0.5 dF/F. Bottom: Heatmaps of all distension-responsive neurons from left (*N* = 4 fish) and right (*N* = 3 fish) vagal ganglia across pulses, sorted by peak response to the first pulse. Color scale indicates dF/F normalized to the baseline of the first pulse (“global dF/F”). (**C**) Percentages of activated (top) and suppressed (bottom) vagal neurons, which were similar between left and right vagus (activated: *P* = 0.2981, suppressed: *P* = 0.3941, unpaired t-tests). Dots represent individual fish. (**D**) Temporal dynamics of activated neurons around injections. Left: average dF/F traces from left (blue, *n* = 31 neurons) and right (orange, *n* = 55 neurons) vagus, normalized to the pre-injection baseline (“local dF/F”). Right: peak local dF/Fs and times to decay showed no significant lateralization (peak dF/F: *P* = 0.6491, Mann-Whitney test, time to decay: *P* = 0.5479, Kolmogorov-Smirnov test). (**E**) Temporal dynamics of suppressed neurons around injections. Peak local dF/Fs and times to decay showed no significant lateralization (peak dF/F: *P* = 0.9001, Mann-Whitney test, time to decay: *P* = 0.7082, Kolmogorov-Smirnov test). (**F**) Average global dF/F traces across five pulses. Darker traces represent increasing distension. (**G**) Peak local dF/Fs and baseline AUCs of activated neurons across pulses. Activated vagal neurons exhibited increasing baseline activity with higher distension (\**P* = 0.0239, mixed-effects analysis and Friedman test). Injection times are marked by red lines (B, representative traces; D–F) and by black lines (B, heatmaps). n.s., not significant. Line and bar graphs show mean ± SEM. Violin plots show median.

Extracting dF/F traces of all vagal neurons, we identified those with stimulus-locked activity (**Figures 3B**, **S4A,** and **S5A**; **Video S3**). Notably, we observed not only neurons activated by gut distension, but also neurons exhibiting suppression (**Figure 3C**). The proportions of activated and suppressed neurons did not differ between left and right vagal ganglia (**Figure 3C**). Similarly, average temporal dynamics of responses to gut distension were comparable between left and right vagal neurons (**Figures 3D** and **3E**), suggesting symmetric encoding of gut mechanosensation in larval zebrafish.

To evaluate how neuronal responses evolve with increasing gut distension, we examined temporal dynamics across consecutive gavage pulses (**Figures 3F**, **3G**, **S5B**, and **S5C**). In both left and right vagal ganglia, activated neurons exhibited progressively elevated baseline fluorescence as distension increased (**Figure 3G**). As population-level analyses can obscure neuron-specific dynamics, we further classified neurons based on their overall peaks across pulses as “integrating” or “habituating” (**Figure S4E**). Only a small proportion of neurons showed integration (left: 7%, right: 10%) or habituation (left: 2.5%, right: 2.3%), with no significant differences between ganglia (**Figure S5D**). This suggests that vagal afferents encode distension predominantly in a monotonic manner, with minimal lateralization between the left and right vagus. These findings support the idea that vagal circuits function as early detectors of distension intensity and reveal that, unlike the lateralized vagal responses observed in mammals^55^, larval zebrafish encode gut distension symmetrically.

### Gut distension broadly activates dorsal hindbrain circuits

To investigate how mechanosensory signals are propagated to downstream brain regions, we next repeated microgavage with water during calcium imaging in the dorsal hindbrain (**Figure 4A**; **Video S4**). Beyond the expected spontaneous or motor-related activity in the hindbrain, we observed fast (<0.8 s), widespread distension-evoked neural activation and suppression in multiple interoceptive regions, including the vagal sensory lobe (VL), area postrema (AP), and vagus motor nucleus (VMN; **Figures 4B–4D**). The anatomical distribution of enterically driven neurons across these regions indicates that distension signals are broadly encoded across the dorsal hindbrain (**Figures 4C** and **S6**). The proportions of activated and suppressed neurons in the hindbrain closely mirrored those in vagal ganglia (**Figure 4D**), suggesting that the responsiveness to gut distension is preserved across these interoceptive nodes and that fast information processing from the gut is already functional in early development.

**Figure 4.**
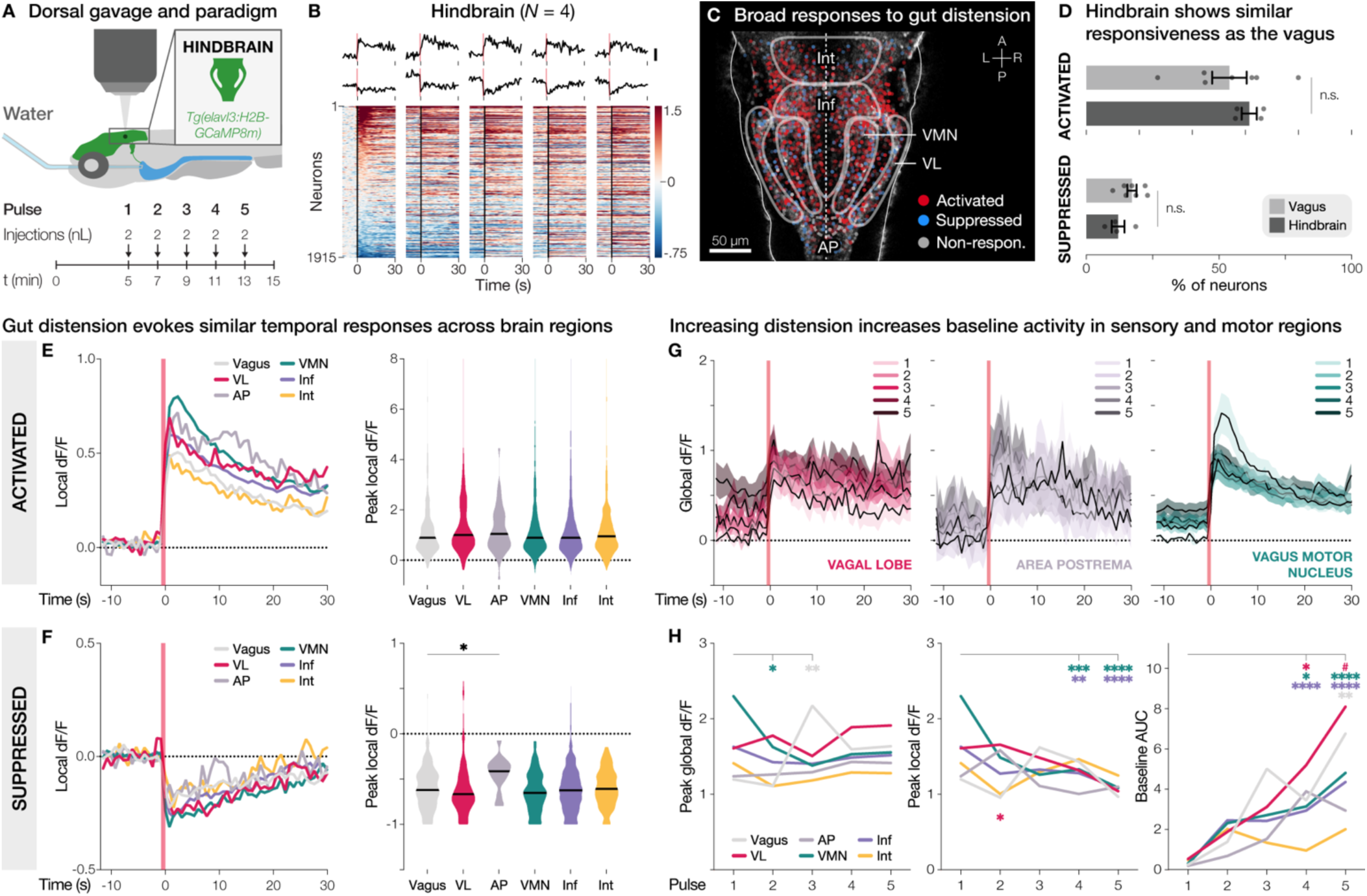
Gut distension evokes widespread neuronal activity across the dorsal hindbrain. (**A**) Experimental setup. 6–10 dpf *Tg(elavl3:H2B-GCaMP8m)* zebrafish were embedded dorsal-side-up and continuously imaged during microgavage with water, delivering five pulses of 2 nL. (**B**) Top: Normalized dF/F traces across five pulses from representative activated (top) and suppressed (bottom) neurons in the hindbrain. Scale bar: 0.5 dF/F. Bottom: Heatmap of all distension-responsive hindbrain neurons (*N* = 4 fish) across pulses, sorted by peak response to the first pulse. Color scale indicates dF/F normalized to the baseline of the first pulse (“global dF/F”). (**C**) Spatial distribution of hindbrain neurons classified as activated (red), suppressed (blue), or non-responsive (gray), plotted on a reference hindbrain. Neurons from all five planes are overlaid. Dot opaqueness indicates higher average peak local dF/F across pulses. Area postrema (AP, dashed line) lies dorsally out of the depicted plane. A: anterior, L: left, P: posterior, R: right. (**D**) Percentages of activated (top) and suppressed (bottom) neurons, which were similar between the vagus and hindbrain (activated: *P* = 0.4259, suppressed: *P* = 0.1106, unpaired t-tests). Dots represent individual fish. (**E**) Temporal dynamics of activated neurons around injections. Left: average dF/F traces from vagus (light gray, *n* = 86 neurons), vagal sensory lobe (VL, pink, *n* = 55 neurons), area postrema (gray, *n* = 8 neurons), vagus motor nucleus (VMN, green, *n* = 291 neurons), inferior dorsal medulla oblongata (Inf, purple, *n* = 879 neurons), and intermediate dorsal medulla oblongata (Int, yellow, *n* = 95 neurons). Right: peak local dF/Fs across regions were not significantly different (*P* = 0.1139, Kruskal-Wallis test). (**F**) Temporal dynamics of suppressed neurons. Left: average dF/F traces from vagus (*n* = 25 neurons), VL (*n* = 21 neurons), AP (*n* = 3 neurons), VMN (*n* = 46 neurons), Inf (*n* = 186 neurons), and Int (*n* = 25 neurons). Right: AP neurons showed less suppression compared to vagal neurons (\**P* = 0.0463, Kruskal-Wallis and Dunn’s multiple comparisons tests). (**G**) Average dF/F traces of activated neurons for pulse 1–5 (light to dark), normalized to the baseline of the first pulse (“global dF/F”). (**H**) Peak global dF/Fs, peak local dF/Fs, and baseline AUCs of activated neurons across pulses. Peak local dF/F: VMN and Inf neurons had smaller changes in peak with increasing distension (*P* < 0.005). Baseline AUC: vagal, VL, VMN, and Inf neurons exhibited increased baseline activity with increasing distension (*P* < 0.03, VL pulse 1 vs. 5: ^#^*P* = 0.0507, mixed-effects analyses with Šídák’s multiple comparisons). Injection times are marked by red lines (B, representative traces; E–H) and by black lines (B, heatmap). \**P* < 0.05, \*\**P* < 0.01, \*\*\**P* < 0.001, \*\*\*\**P* < 0.0001. n.s., not significant. Line and bar graphs show mean ± SEM. Violin plots show median.

The average temporal dynamics of activated neurons were also conserved across vagal and hindbrain regions (**Figure 4E**). Suppressed neurons also displayed similar response profiles, except for AP neurons, which exhibited significantly weaker suppression compared to vagal neurons (**Figure 4F**). To examine hindbrain responses to increasing gut distension, we compared neural activity across sequential pulses (**Figure 4G**). VMN neurons showed progressively smaller peaks with higher distension intensities, suggesting adaptation (**Figure 4H**). Notably, baseline fluorescence increased over repeated pulses in the VL, VMN, and inferior dorsal medulla oblongata (Inf), mirroring trends observed in vagal neurons (**Figure 4H**). These results indicate that while immediate responses in motor regions diminish with increasing distension, baseline activity steadily rises in both sensory and motor areas. This broad and fast encoding across the dorsal hindbrain highlights gut distension as a salient interoceptive signal in larval zebrafish.

### Neuronal response subtypes to increasing gut distension

To characterize neural responses across repeated pulses, we categorized activated and suppressed neurons as “monotonic”, “integrating”, or “habituating” based on their peak amplitudes (**Figure S7A**). Among these, activated & monotonic, suppressed & monotonic, and activated & integrating subtypes were consistently observed across both the vagus and hindbrain in all fish (**Figures S7B** and **S7C**). The proportions of these subtypes did not differ significantly across regions (**Figure S7D**), and subtypes appeared not to be spatially clustered (**Figure S8**), suggesting distributed recruitment rather than regional specialization for response dynamics to distension.

We next characterized the temporal profiles of each subtype across regions (**Figure S9**). While monotonic neurons were defined by the absence of significant trends in peak dF/Fs, activated & monotonic neurons in the VMN showed diminishing peaks and shorter decay times over successive pulses, resembling habituation-like dynamics (**Figures S9A** and **S9B**). In contrast, activated & integrating neurons exhibited consistent increases of peak responses across pulses in all regions, reflecting progressive integration of mechanosensory input (**Figures S9E** and **S9F**). Interestingly, baseline fluorescence rose across all subtypes in the VMN and Inf, potentially reflecting global increases in neural activity driven by slow hormonal feedback (**Figure S9**). Together, these findings reveal diverse and distributed spatiotemporal coding strategies for encoding enteric distension signals across vagal and hindbrain circuits in early development.

### Chemosensory encoding of AITC features slow-onset dynamics

To examine how additional chemosensory information influences gut-brain signaling, we injected nutrient solutions at the same concentration as in the CPs of feeding experiments (25 mM; **Figure 1**). Based on findings from our gut-mediated feeding assay and previous literature^63,86^ (**Figure 1F**), we selected glucose as a potentially neutral, glycine as an appetitive, and AITC as an aversive chemical stimulus (**Figure 5A**). Repeating our microgavage protocol of five pulses of 2 nL with these nutrients allowed us to compare across neurons in the vagus and dorsal hindbrain (**Figure 5B**). As with water microgavage, we observed almost no lateralization between the left and right vagus across nutrient stimuli (**Figure S10**). Given the overall symmetry of vagal responses, data from both ganglia were combined for subsequent analyses.

**Figure 5.**
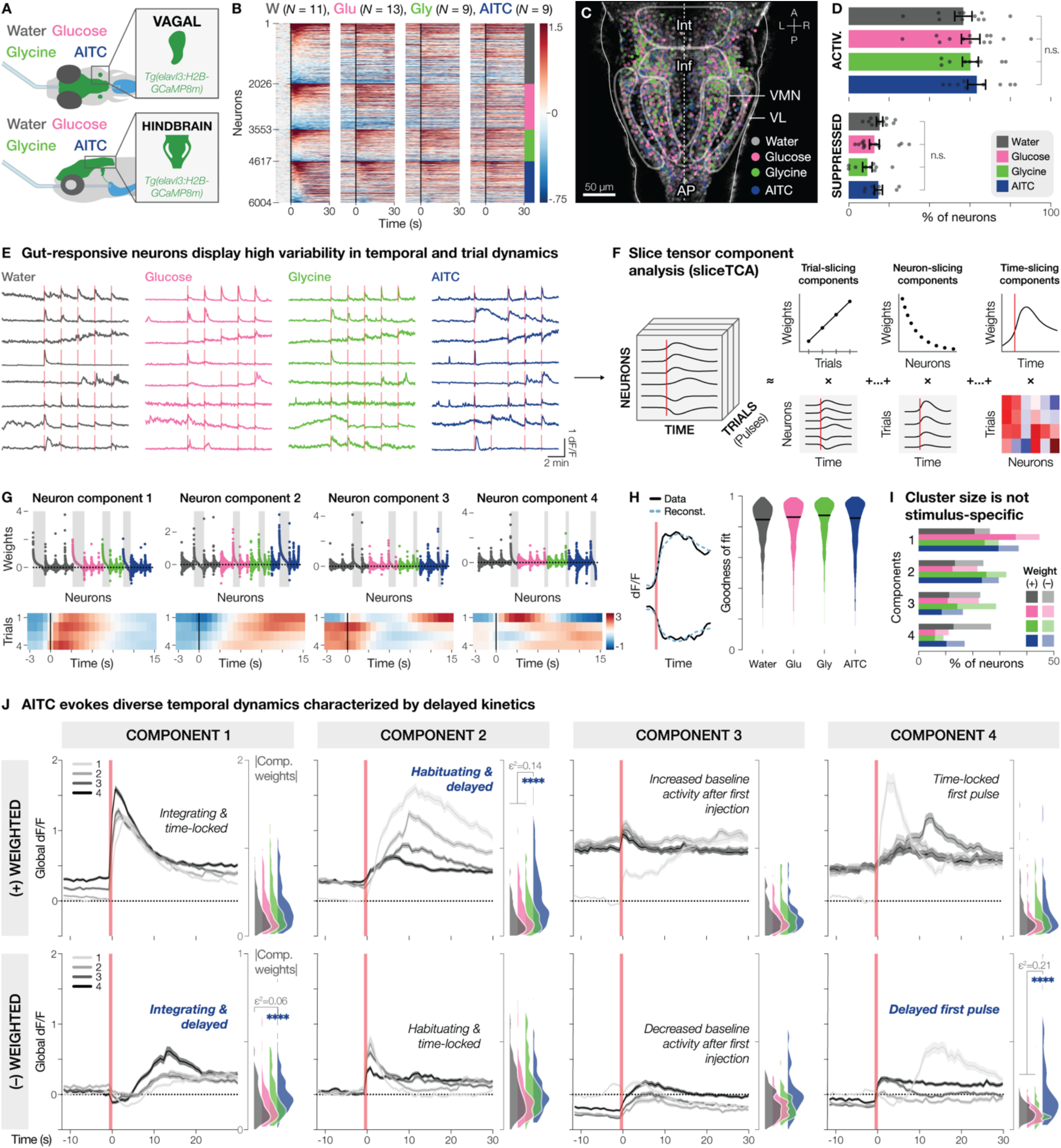
Enteric stimuli drive complex temporal dynamics in vagal and hindbrain neurons. (**A**) Schematic of imaging configurations to record gut-evoked neural activity in vagal and hindbrain neurons in response to microgavage of water, glucose (25 mM), glycine (25 mM), or AITC (25 mM). (**B**) Heatmap of all stimulus-responsive vagal and hindbrain neurons across pulses, sorted by peak response to the first pulse (water: *N* = 11 fish, gray, glucose: *N* = 13 fish, pink, glycine: *N* = 9 fish, green, AITC: *N* = 9 fish, blue). Black lines indicate injection times. Color scale indicates global dF/F. (**C**) Spatial distribution of gut-responsive hindbrain neurons colored by stimulus identity. Neurons from all five planes are overlaid. Dot opaqueness indicates higher average peak local dF/F. Brain regions as in **Figure 4C**. A: anterior, L: left, P: posterior, R: right. (**D**) Percentages of activated (top) or suppressed (bottom) neurons were not significantly different across stimuli (activated: *P* = 0.7819, one-way ANOVA, suppressed: *P* = 0.1747, Kruskal-Wallis test). Dots represent individual fish. (**E**) Representative normalized dF/F traces for all enteric stimuli showing temporal and trial variability. (**F**) Schematic of slice tensor component analysis (sliceTCA). Traces from all stimuli and regions were compiled into a 3D data tensor (trials x neurons x time). The first four successful trials were used for consistency. SliceTCA decomposes this tensor into trial-, neuron-, and time-slicing components, where the data tensor is approximated as the outer product of each component’s weights and slice. (**G**) Neuron-slicing components of the sliceTCA model. Weights are colored by stimulus. Neurons were clustered into the component with their highest weight (gray bars). Note that neurons may have positive or negative weights for their assigned component. Black lines indicate injection times. Color scale indicates slice value. (**H**) Left: representative trace reconstructions (blue dashed line) overlaid on raw data (black). Right: reconstruction goodness of fit per neuron, grouped by stimulus. Black lines show median. (**I**) All enteric stimuli drove all four temporal neural dynamics, illustrated by the percentage of positive (darker) and negative (lighter) weighted clusters per stimulus, averaged across fish without significant differences across stimuli (mixed-effects analysis and Šídák’s multiple comparisons test). (**J**) Left: average dF/F traces of positive- and negative-weighted neurons per component, plotted across pulses (light to dark), increasing gut distension. Right: Stacked violin plots show absolute component weights per stimulus. Clusters C1–, C2+, and C4– had significantly stronger weights in AITC-responsive neurons, indicating AITC evokes diverse delayed dynamics. Other clusters had no significant differences across nutrients, reflecting the importance of distension-related dynamics. Only comparisons with *P* < 0.05 (Kruskal-Wallis and Dunn’s multiple comparisons tests) and at least medium effect sizes (ε^2^ ≥ 0.06 with epsilon-squared and |δ| ≥ 0.33 with Cliff’s delta) were considered significant. Injection times are marked by red lines (D, E, I, and J). \*\*\*\**P* < 0.0001. n.s., not significant. Mean ± SEM.

Next, we assessed how chemosensory signals were encoded across the dorsal hindbrain (**Figures 5C**, **S11A**, and **S11B**). Similar proportions of activated and suppressed neurons were observed between the vagus and hindbrain and across stimuli (**Figures 5D** and **S11C**). However, spatial analyses revealed stimulus-specific topographies in the hindbrain, with more anteriorly distributed glycine-responsive neurons and more posterior AITC-responsive neurons (**Figure S12A**). This separation was significantly pronounced in the VL, where glycine- and AITC-responsive neurons showed the greatest spatial divergence (**Figure S12B**). Average temporal dynamics revealed that AITC evoked significantly slower activation in VL, VMN, and Inf neurons (**Figures S11D** and **S11E**). These results suggest that AITC drives distinct temporal activity within the vagus-hindbrain circuit, supporting the notion that the brain differentially encodes gut-derived aversive signals through stimulus-specific temporal dynamics.

### Enteric information is encoded in diverse temporal dynamics

To further dissect the neural dynamics evoked by repeated gut stimulation, we analyzed temporal patterns across pulses for all responsive vagal and hindbrain neurons (**Figure 5E**). Prompted by the heterogeneity in neuronal responses across time and injection pulses, we employed the unsupervised dimensionality reduction method slice tensor component analysis^87^ (sliceTCA; **Figure 5F**). SliceTCA decomposes a three-dimensional data tensor, which in our case is comprised of the injection pulses, neurons, and time, into separate components along each dimension, thus enabling a biologically interpretable representation of diverse response patterns.

An optimized parameter search that balanced cross-validation loss, reconstruction performance, and neuron-weight sparsity yielded a model with 0 trial-slicing, 4 neuron-slicing, and 1 time-slicing component **(Figures 5G, 5H**, and **S13**). Each of the four neuron-slicing components (C1–4) revealed pairs of distinct temporal dynamics, resulting in eight total neuron clusters. We assigned each neuron to one of these eight clusters based on its sign (positive (+) or negative (–)) and the magnitude of its strongest neuron-slicing component weight. These clusters were similarly represented across the left and right vagus, brain regions, and nutrient stimuli (**Figure 5I**; **Table S2**), indicating the anatomically widespread recruitment of these temporal motifs, regardless of stimulus type or brain region.

Temporal analysis of component dynamics revealed distinct biological response signatures. While C1 captured integrating neurons across increasing gut distension (water: 26.46%, glucose: 44.85%, glycine: 29.73%, AITC: 37.05%), C2 clustered habituating neurons (water: 23.82%, glucose: 21.71%, glycine: 32.33%, AITC: 29.47%). Interestingly, C3 and C4 reflected unique dynamics associated with the onset of sensory input. C3 was defined by changes in baseline fluorescence after the first injection (water: 22.92%, glucose: 22.39%, glycine: 28.78%, AITC: 16.48%), whereas C4 characterized neurons that responded strongly to the initial pulse (water: 26.80%, glucose: 11.05%, glycine: 9.16%, AITC: 17.00%) (**Figures 5J** and **S14**). While components C1 and C2 dynamically encoded the ongoing state of gut content, components C3 and C4 likely represent the initial sensory enteric detection, potentially signaling confirmation of consumption to the brain.

Finally, we assessed the contributions of each stimulus-responsive neuron to individual clusters. Most clusters exhibited comparable component weights across all nutrient stimuli as well as water, reinforcing that gut distension is the dominant visceral cue in larval zebrafish, even in the presence of nutrients that modulate feeding behavior (**Figure 1F**) and are known to activate EECs^62,63^. However, three specific clusters (C1–, C2+, and C4–) displayed significantly higher weights from AITC-responsive neurons (**Figure 5J**). All three clusters exhibited slower response onsets with prolonged activity, indicating that AITC consistently drives slower activation patterns. These findings demonstrate that while gut distension initiates robust neural responses with fast onsets, AITC drives temporally distinct, slow-onset populations across the vagus and hindbrain.

## Discussion

Understanding how post-ingestive signals from the gut are communicated to the brain is critical for identifying the neural basis of feeding behaviors. By measuring the consumption of CPs that bypass exteroceptive chemical signaling, we found that zebrafish EECs regulate feeding behavior during early development. By developing a novel microgavage method to deliver enteric stimuli during calcium imaging, we demonstrate that rapid gut-brain communication is present as soon as general gut function develops. Specifically, we show that zebrafish vagal afferents and their hindbrain target neurons encode diverse enteric signals, exhibiting rich temporal dynamics of activation and suppression related to feeding status and history. These results extend our understanding of interoceptive signaling, revealing that gut-derived mechanical and chemical stimuli modulate brain activity in less than a second of ingestion, even in the developing vertebrate brain (**Figure 6A**).

**Figure 6.**
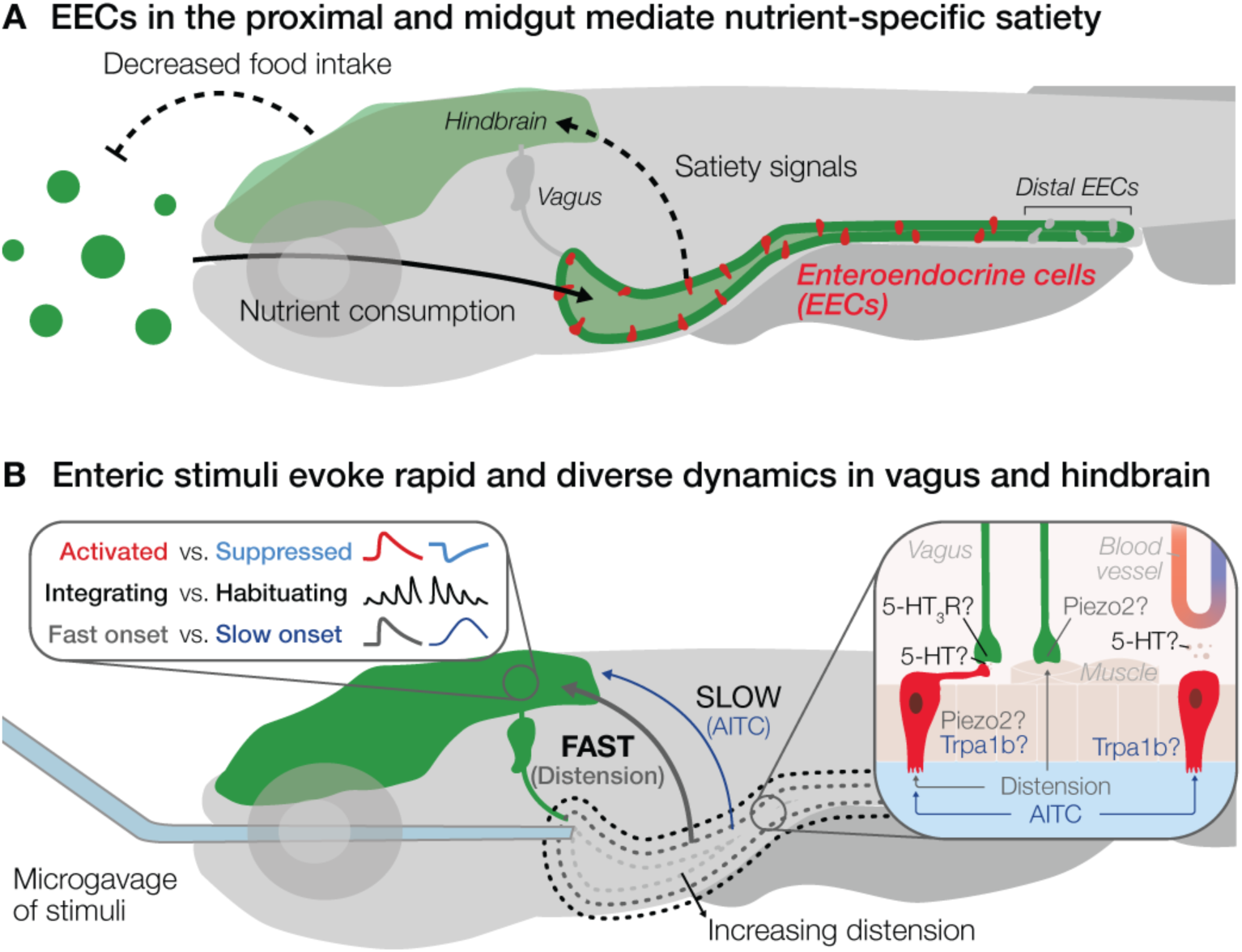
Graphic summary of enteric information processing in larval zebrafish. (**A**) Cartoon illustrating the role of EECs in the proximal and midgut in regulating food approach and post-ingestive feeding. Upon nutrient consumption, like glucose or glycine, EECs communicate satiety signals that act on the brain to decrease food intake. (**B**) Enteric stimuli evoke diverse temporal dynamics in vagal and hindbrain neurons. While gut distension is encoded with fast, time-locked neural signatures, the aversive compound allyl isothiocyanate (AITC) induces responses with prolonged peaks. While distension may activate Piezo2 channels on neuropod-forming EECs and/or Piezo2*-*positive vagal afferents that innervate the gut musculature, AITC activates Trpa1b on EECs^63^, which may signal enteric information synaptically and/or hormonally. Both Piezo2 and Trpa1b are expressed in serotonin-producing enterochromaffin cells, suggesting that serotonin signaling via 5-HT3 receptors on vagal neurons may contribute to gut-brain communication.

To investigate whether and how enteric signaling contributes to feeding in larval zebrafish, we developed a gut-mediated feeding assay using CPs, which have been used in aquaculture to enhance nutrient retention in marine feeds^71,74^. Adapting this approach for freshwater zebrafish larvae enabled us to isolate the gut-mediated post-ingestive effects of appetitive and aversive nutrients on feeding. Glycine and alanine, amino acids abundant in the zebrafish’s natural diet^82,88–90^, robustly promoted feeding. In contrast, glucose, which drives consumption in mammals^91^, had no such effect in larval zebrafish, despite high concentrations (500 mM) of glucose activate EECs in zebrafish^62^. This nutrient-specific behavioral preference aligns with ecological observations, as zebrafish prey contains high levels of glycine, potentially serving as a chemical cue for foraging^81,82,92^. Additionally, the zebrafish homologue of the mammalian sweet taste receptor is more responsive to amino acids than sugars^93^, reinforcing the relevance of amino acids over glucose in zebrafish feeding.

By comparing feeding in wild-type and zebrafish with depleted EECs^63^, we uncovered a dual role for EECs, promoting food approach, yet regulating the consumption of specific nutrients once feeding was initiated. This dichotomy suggests a functional division among EEC subtypes^66^. Indeed, ghrelin-expressing EECs likely promote hunger and food approach^66,94,95^, while others release satiety peptides, like cholecystokinin and peptide YY, to limit intake once nutrients are ingested^5,33,62,66,96,97^. Removing these regulatory EECs may thus disrupt the balance between motivation to feed and satiation, resulting in increased intake after feeding has occurred. These findings highlight the multifaceted roles of EECs in shaping both anticipatory and post-consummatory aspects of feeding behavior shortly after the onset of exogenous feeding^60,61^.

To begin dissecting how post-consummatory gut signals are relayed to the embryonic brain, we developed a novel microgavage^85^ method to deliver specific nutrients into the small gut lumen while simultaneously recording neural activity at single-cell resolution. Importantly, this approach allows us to mimic naturalistic feeding by activating both mechanosensory and chemosensory signaling in awake zebrafish. In less than a second after enteric stimulus delivery, we observed widespread activation and suppression across vagal and hindbrain neurons. Our results revealed rapid gut-brain communication as early as 6 dpf, soon after the formation of a functional gut, suggesting the presence of fast synaptic mechanisms in early development^1,9^.

Since the gut inherently distends during nutritive stimulus delivery, we first focused on mechanosensory signals. A key candidate for mediating distension detection is the mechanosensitive ion channel Piezo2, which has been implicated in sensing distension in peripheral organs, including the gastrointestinal tract^98–100^, and is conserved from mammals to fish^101–103^. Notably, both EECs and vagal afferents express Piezo2^67,68^, suggesting that mechanical signals may be sensed in multiple sensors along the gut-brain circuit^100,104,105^ (**Figure 6B**). In mammalian EECs, Piezo2 is enriched in enterochromaffin cells and can initiate serotonin release in response to distension, modulating downstream neural activity^31,106,107^. Alternatively, vagal afferents expressing Piezo2 can directly respond to mechanical forces, enabling rapid neural signaling^105^. The co-expression of Piezo2 in both cell types suggests a multilayered mechanism for encoding gut distension. Future studies should aim to disentangle EEC-derived mechanosensation from direct activation of vagal afferents.

With access to the entire vagal population in larval zebrafish, we observed minimal response lateralization to distension, contrasting with the functional asymmetries of the left and right vagus observed in rodents^6,108,109^. This discrepancy may reflect underlying anatomical differences. Compared to the asymmetric gastrointestinal tract of adult mammals and lateralized vagal innervation and function, the larval zebrafish gut is largely symmetrical^110,111^, which may underlie the absence of functional lateralization in zebrafish. Investigating how gut anatomy and vagal function develop may reveal whether lateralized gut-brain signaling emerges in conjunction with bodily asymmetry.

A key yet underexplored feature of gut-brain communication is the temporal coding of visceral signals. By delivering repeated intraluminal stimuli that mimicked multiple gulps or bites, we uncovered a diversity of temporal dynamics in neural responses across vagal and hindbrain neurons. These response patterns demonstrate that enteric information about ongoing feeding status and history is relayed in neurons exhibiting sensitization, habituation, delayed onset, ramping, and selective responses to initial trials, revealing how neural dynamics evolve on a trial-by-trial basis (**Figure 6B**). Our data emphasize that interoceptive coding is inherently dynamic, and that uncovering the gut’s influence on the brain requires attention to how signals unfold across time and experience.

Interestingly, enteric stimulation elicited not only activation but also suppression in subsets of vagal afferents and dorsal hindbrain neurons. While excitatory vagal responses to enteric stimuli are well documented^12,14,15,50,52,53,55,112^, suppressive dynamics have also been described across visceral sensory systems, particularly in classical electrophysiological studies and under specific physiological conditions. Early work in mammalian vagal afferents identified slow, calcium-dependent afterhyperpolarizations in subsets of sensory neurons, producing prolonged suppression following brief stimulation and shaping excitability over seconds-long timescales^113–117^. Such intrinsic membrane properties provide a plausible cellular substrate for the suppressive signals observed here. In parallel, inhibitory modulation may arise from neurotransmitter-mediated mechanisms. GABA-positive enteroendocrine cells have been reported in the gastrointestinal tract in early histochemical studies^118–120^, and vagal sensory neurons express both ionotropic and metabotropic GABA receptors^53,121^, which have been shown to gate mechanosensory responsiveness in mammalian systems^122–125^. These peripheral inhibitory pathways may therefore act alongside excitatory glutamatergic inputs to balance vagal signaling and sharpen contrast between competing visceral signals. Future studies using neurotransmitter-specific reporters or pharmacological tools could clarify the role of suppression in interoceptive coding.

Here, we show that in awake larval zebrafish, gut distension alone is sufficient to elicit rapid and widespread neural responses across vagal and dorsal hindbrain neurons. Our findings suggest that increasing gut distension drives distributed neural activity, complementing a recent elegant study in larval zebrafish that excluded mechanical cues by local photouncaging glutamate and glucose in the gut, which reported more spatially restricted responses in the hindbrain^70^. This ubiquity of distension-responsive neurons here underscores gut distension as a dominant visceral signal for fast neural communication during this developmental window.

The distributed neural responses observed here may also reflect the nature of the stimulation paradigm. Many studies reporting spatially organized visceral representations have relied on highly focal activation approaches. For example, focal ultraviolet activation of Trpa1-positive vagal neurons spanning the anterior-posterior pharyngeal arches produced corresponding topographic response patterns along the hindbrain axis^73^, and spatially restricted uncaging of nutrients in the gut elicited localized activity within the dorsal vagal complex and dorsomedial hindbrain^70^. In contrast, our microgavage approach more closely mimics naturalistic gut distension by inducing volumetric mechanical stimulation across extended regions of the intestine. This method engages not only the proximal gut but, at higher volumes, also the mid- and distal intestine, likely recruiting multiple mechanosensory pathways simultaneously that converge onto overlapping central circuits. Such broad, multimodal activation is therefore expected to reduce fine-grained spatial segregation and instead produce the distributed hindbrain response patterns observed in awake animals.

In our study, glucose and glycine in the gut evoked neural responses that were nearly indistinguishable from those evoked by distension alone. While these results may be specific to the conditions tested, such as volume or concentration of delivered stimuli, these same concentrations were sufficient to elicit behavioral changes in our gut-mediated CP feeding assay. Therefore, it is possible that nutritive information is overpowered by the dominant stretch signal shared across all stimuli. Systematically varying nutrient concentrations and decoupling mechanical from chemical cues will be essential to clarify how distension and nutrient identity interact to influence feeding-related brain activity.

Nonetheless, the aversive compound AITC triggered diverse neural dynamics with a unifying characteristic of slow-onset kinetics. This slower temporal profile mimics the dynamics of visceral pain signaling, which has a slower onset than somatic pain^128–130^. Zebrafish also use earlier sensory modalities such as olfaction^77,131,132^, taste^93,133–135^, and the lateral line^134,136^ to avoid harmful substances pre-ingestion. Slow-onset enteric signals may therefore function as a failsafe, activating post-ingestive defenses like excretion, emesis, or feeding cessation when earlier checkpoints fail to detect the threat.

Our results highlight that in early development, EECs in the gut epithelium already begin to shape feeding behavior by mediating nutrient-specific satiety signals to the brain. Among these post-ingestive signals transmitted via the gut-vagus-brain circuit, we found that rapid detection of gut volume changes and aversive stimuli takes precedence over identifying specific nutrient content. This prioritization likely reflects the need to rapidly assess how much has been ingested to meet the high and variable energy demands of early development^137,138^, when individual gut capacities and intake can differ significantly^139–141^. As animals mature and their feeding strategies become more selective, we expect finer sensory discrimination mechanisms to gain prominence. While fast gut-brain communication is established in adult systems, our findings demonstrate that this essential survival strategy for evaluating ingestion begins far earlier in life than previously appreciated.

## Resource availability

### Lead contact

Further information and requests for resources and reagents should be directed to the lead contact, Eva A. Naumann.

### Materials availability

All the original reagents presented in this study are available from the lead contact on a reasonable request.

### Data and code availability

All data reported in this paper will be shared by the lead contact upon request. All analyses were run on custom Python scripts, Jupyter notebooks, and GraphPad Prism files, available on https://gitlab.oit.duke.edu/ean26/arinel-et-al_2025. For calcium imaging analysis, the notebooks utilize our custom-written module at https://github.com/Naumann-Lab/caImageAnalysis/tree/arinel-et-al_2025. The notebooks for alignment use another environment and repository, which can be found at https://github.com/Naumann-Lab/alignment. Detailed instructions for executing the analysis notebooks and replicating published results are provided in our manuscript repository.

## Acknowledgements

We thank J. Rawls, L. Ye, and M. Morash for providing transgenic lines, training in the original gavage method, and insightful discussions and feedback; A. Oesterle for assistance in optimizing gavage needle production; M. Bagnat for microforge access; A. Pellegrino for assistance with sliceTCA optimization and calculations related to goodness-of-fit and sparsity; D. Bohórquez for insightful discussions on gut-brain sensory biology; and K. Abdelaal for feedback on experiments, figures, and the manuscript. We would also like to thank J. Burris, K. Olivera, and L. Frauen for zebrafish husbandry. This work was supported by a Boehringer Ingelheim Fonds PhD fellowship to M.A., by a National Science Foundation Graduate Fellowship to K.M.M., as well as a Duke Institute of Brain Sciences Research Incubator Award and an Alfred P. Sloan Foundation Research Fellowship to E.A.N.

## Author contributions

M.A. and E.A.N. conceived the project. M.A. validated the novel transgenic lines and was responsible for conducting and analyzing all gavage experiments. M.A. also developed the gavage methodology, with assistance from K.M.M.. J.A. conducted the gut-mediated feeding experiments and handled the crossing of zebrafish lines for all experiments. E.P.D. generated the new transgenic zebrafish lines. M.H. created all the complex particles used in the feeding experiments. M.A. and E.A.N. wrote the manuscript and prepared the figures with feedback from all contributing authors.

## Experimental model and study participant details

### Animals

For all experiments, we used 6–10 dpf zebrafish as their gastrointestinal tract is functional^142^, and the yolk sac is depleted^143^ at this age. Zebrafish are obtained as fertilized embryos from adult breeders, maintained at a 14/10-hour light/dark cycle at 28.5°C. Groups of ∼30 embryos were raised in pH-buffered, filtered E3 medium (5 mM NaCl, 0.55 mM CaSO_4_, 0.45 mM NaHCO_3_). Starting at 4 dpf, the zebrafish were fed daily with live paramecia. For feeding behavior experiments, we used Casper zebrafish, which lack dark melanophores and reflective iridophores on the skin^144^. We tested the effects of EEC depletion on feeding in outcrosses of *Tg(neurod1:cre; cmlc2:EGFP)^rdu^*^79^ and *TgBAC(gata5:loxP-mCherry-STOP-loxP-DTA)^pd^*^315^ EK (Ekwil) zebrafish, which express diphtheria toxin (DTA) in most of the EECs located in the proximal and midgut^63^. For anatomical imaging of EECs and vagal fibers (**Figure 2A**), we outcrossed *Tg(−5kbneurod1:TagRFP)^w^*^69145^ to *Tg(isl1:EGFP)^rw^*^0146^. For two-photon imaging in vagal neurons and hindbrain, we generated two novel transgenic lines, *Tg(elavl3:H2B-GCaMP8m)* in the Casper background with nuclear-localized expression of the calcium sensor GCaMP8m *and Tg(neurod1:GCaMP8m-SV40)* with standard molecular cloning techniques (**Supplementary methods**). All experiments were approved by Duke University School of Medicine’s Institutional Animal Care and Use Committee (IACUC, Protocol Registry Number A058-24-03).

## Method details

### Gut-mediated feeding assay

To test gut-mediated feeding, we developed a behavioral assay using complex particles (CPs), which are odorless, alginate-encapsulated liposomes that release nutrients post-ingestion. CPs were produced as previously described^74,147^ in the 50–150 µm size range (**Supplementary methods**). We separated 8-dpf sibling zebrafish into new petri dishes of control and nutrient groups (maximum 20 larvae per dish), each containing 55 mL of E3 medium. After 24 hours of fasting, we introduced 18 mL of CP solution into each dish for a final concentration of 0.05 mg/mL. Control particles contained only sodium fluorescein as a negative control. Test particles contained sodium chloride (NaCl), D-(+)-glucose, glycine, or L-alanine (**Table S1**). Particle solutions were prepared at a concentration of 0.2 mg/mL in E3 medium and stored at 4°C and warmed to room temperature 30 minutes before each experiment. CP solutions were filtered using a 70 µm strainer (76327-100, VWR) to remove large particles. Three experiments were conducted without filtering but showed no significant effect on feeding (see “Feeding behavior data processing and analysis”). After free feeding for 45 minutes, zebrafish were transferred into 50 mL Falcon tubes containing 20 mL E3 medium and placed on ice. Post-euthanasia, E3 medium was replaced with 10 mL of 4% (w/v) paraformaldehyde (PFA; 047340.9M, Thermo Scientific) in 1X phosphate-buffered saline (119-069-101, Quality Biological) for fixation. After an hour, the fish were moved to 15 mL Falcon tubes containing 10 mL of 0.1% Triton X-100 (9002-93-1, MP Biomedicals) in E3 medium for repeated washing. Individual zebrafish were either immediately embedded on their side in 1.5% (w/v) low-melting-point agarose (16520-050, Invitrogen) or stored overnight at 4°C for next-day imaging. Side-view images were captured using a stereo microscope (MVX10, Olympus) at 6.3X zoom with blue light illumination (X-Cite mini+ Compact, Excelitas) and a smartphone (iPhone SE, Apple) with a phone-to-eyepiece adapter (Gosky, https://www.amazon.com/gp/product/B013D2ULO6). All images were recorded with the ProCam app^148^, setting shutter speed to 1/10 seconds, ISO at 2112, and white balance at 4400K. Images were saved directly as TIFF files at resolutions of 3024×3024 pixels for square images and 4032×3024 for 4:3 ratio images.

### Feeding behavior data processing and analysis

After acquisition, images were imported to Fiji as separate TIFF files. For each fish, we selected the entire gut area, avoiding autofluorescence from the swim bladder, particles in the esophagus that have not reached the gut, or debris outside the fish. The cropped gut images were saved as separate 8-bit grayscale TIFF files. Applying a conservative threshold to extract only the particles, we used custom Python code to analyze fluorescence intensity values per fish. To assess whether CP filtration affected the results, we performed a bootstrapping analysis comparing the gut fluorescence of controls of filtered and unfiltered experiments. Across 1000 iterations, we found no significant difference between filtered and unfiltered conditions (median *P* = 0.3937, Kolmogorov-Smirnov test), so we combined all datasets. Only fish with fluorescence in their gut were included in pixel intensity analyses, reflecting active feeding behavior. To determine batch variability in consumption, we compared raw fluorescence values of control particles across experiments (**Figure S1B and S1D**), which showed significant batch effects (*P* < 0.0001, Kruskal-Wallis test). Therefore, we normalized raw pixel intensity values to the median of each experiment’s control distribution.

### Microgavage during two-photon imaging

#### Microgavage needle preparation

To record neural activity in response to enteric stimulation, we adapted the zebrafish microgavage procedure^85^ to deliver precise amounts of nutrients to the intestinal lumen of awake zebrafish during two-photon imaging. To prepare custom microgavage needles, we used a pipette puller (P-1000, Sutter Instrument) with 1.0 mm aluminosilicate glass (AF100-64-10, Sutter Instrument), which is stronger and more flexible than borosilicate^149^. Needles were clipped to an outer diameter of ≤20 µm using the glass-on-glass technique^149^, bent 30° 5 mm from the tip, and lightly fire-polished using a microforge (MF2, Narishige). On each experimental day, individual gavage needles were back-filled with mineral oil (M5904, Sigma-Aldrich), using a 1 cc syringe (748019-0001, Kimble) connected to a pipette filler (MF28G-5, World Precision Instruments), removing any air bubbles. Once filled, each needle was attached to a holder, which is linked directly to a separate syringe barrel (910-N, Metcal), itself connected to an injector (DX-250, Metcal). The needle and syringe assembly was then mounted on a manual micromanipulator (M-152, Narishige). 1 µL of phenol red (P0290, Sigma-Aldrich) was mixed with 4 µL of the selected stimulus on parafilm. <1 µL of this solution was then front-filled into the needle. To calibrate the injection volume, the injector air pressure and time were adjusted to result in consistent 2 nL ejections without any capillary action (**Supplementary methods**).

#### Stimulus preparation for gavage

For stretch-only experiments, gavage needles were filled with E3 medium. For chemical stimuli, 31.25 mM solutions were prepared and mixed 1:4 with phenol red (P0290, Sigma-Aldrich) to yield a final concentration of 25 mM. D-(+)-glucose (G7021, Sigma-Aldrich) and glycine (G7126, Sigma-Aldrich) were dissolved in E3 medium, while allyl isothiocyanate (AITC; 377430, Sigma-Aldrich) was prepared in 1X phosphate-buffered saline (70011-044, Gibco) to ensure solubility^63^.

#### Microgavage during calcium imaging

We selected 6–10 dpf *Tg(elavl3:H2B-GCaMP8m)* zebrafish with bright expression. Depending on the experiment, we embedded individuals either sideways (vagal imaging) or dorsal side up (hindbrain imaging) in the center of a 55 mm petri dish’s lid, onto a layer of 1.5% (w/v) low-melting-point agarose (16520-050, Invitrogen). After flooding the dish with E3 medium, a fine scalpel (08-924-20, FEATHER 715) removed agarose around the fish’s mouth. We used a larger scalpel (MPR-47101, MedPride) to cleanly trim agarose on both sides of the fish, ensuring the proper position of a right-angle prism mirror with anti-reflective coating on its legs (PS910L-B, Thorlabs) and minimal obstruction for the infrared light. After positioning of the fish under the two-photon objective, the syringe barrel with the gavage needle was mounted onto a motorized micromanipulator (MP-865, Sutter Instrument) on a custom acrylic stage. With an infrared light (M780L3, Thorlabs) directed at the prism, we captured images through a 45° hot mirror (43-958, Edmund Optics) reflecting light towards an infrared camera (1800 U-2050m, Allied Vision). Guided by the dual view of anatomical landmarks, including the mouth, heart, swim bladder, otolith, gut, and pharynx, the gavage needle was inserted into the mouth without harming the fish. Using fine control of the micromanipulator, the needle was gently moved past the esophagus into the proximal gut, monitoring the fish’s heartbeat and watching for signs of twitching, indicating potential injury. Each experiment started by recording a 5-minute baseline calcium activity in vagal or hindbrain populations, followed by five injections administered at 2-minute intervals, and an additional 2-minute recording. After each gavage, we slowly retracted the needle using the micromanipulator. Afterwards, we examined each fish under a stereo microscope to confirm that the phenol red is localized only within the gut. Checking for normal heart rate, liver morphology, and jaw integrity, fish were released from the agarose to assess their motor behavior.

### Two-photon calcium imaging

To record neural activity in live zebrafish, we used a two-photon laser-scanning microscope (Ultima2Pplus, Bruker) with a 20X objective (XLUMPLFLN20XW, Olympus). Fluorescent images were captured using a pulsed femtosecond laser (Axon 920-2 TPC, Coherent) as excitation source, at a power of 23 mW on the specimen. For functional volumetric recordings, we imaged at a resolution of 512×512 pixels, covering 200.5 µm x 200.5 µm for vagal and 267.3 µm x 267.3 µm for brain recordings. We averaged 4 frames per acquisition, achieving a final rate of ∼1.3 Hz. Using resonant scanning, we recorded 5 planes volumetrically, spaced 10 µm apart, in either left or right vagal neurons or hindbrain, identified using anatomical landmarks in the *Tg(elavl3:H2B-GCaMP8m)* line. At the end of each imaging session, we captured a high-resolution (1024×1024 pixels) anatomy stack from 10 µm above to 10 µm below the targeted imaging planes, at intervals of 2.5 µm. Left and right vagal ganglion imaging was performed in separate fish due to anatomical separation.

### Two-photon data processing and analysis

Following data acquisition, we preprocessed and aligned uncompressed image volumes with their corresponding timestamps for gavage injections, for analysis of individual planes. Using CaImAn^150^, we performed motion correction with NoRMCorre^151^ and source extraction with constrained non-negative matrix factorization (CNMF)^152^. Parameters were optimized via mesmerize-core (https://github.com/nel-lab/mesmerize-core). The resulting sources, referred to as neurons, were filtered based on criteria such as location, temporal peaks, and proximity to other neurons. To observe spatial organization, we aligned each recording to an anatomical reference image using reverse alignment with SimpleITK^153–155^.

All heatmaps display denoised traces, sorted by the onset of its maximum response. From these data, we characterized five temporal characteristics: (1) peak dF/F normalized to the pre-injection baseline of the first trial (global dF/F), (2) peak dF/F normalized to the pre-injection period of each trial (local dF/F), (3) baseline area under the curve (AUC), (4) time at half-maximum, and (5) time to decay (**Figures S4C** and **S4D**, **Supplementary methods**). We used Spearman rank-order correlation on the peak global dF/F values to categorize neurons as “integrating” or “habituating” (*P*<0.05) (**Figure S4E**). Remaining non-significant neurons were categorized as “monotonic”. We focused on subtypes and regions with at least 10 neurons total for vagal recordings or 10 neurons per fish for hindbrain regions (**Figure S7**).

### Slice tensor component analysis (sliceTCA)

#### Data preprocessing

To focus sliceTCA dimensionality reduction on the post-injection period, we extracted traces from 4 frames before and 20 after each injection of responsive neurons. Since some zebrafish only received four injections, we focused on the first four pulses. All traces were smoothed with a Gaussian filter along the time axis (σ=1) and normalized to the overall standard deviation so that the mean squared error corresponded to the unexplained variance.

#### Cross-validated model selection

To choose the number of components in each slice type, we ran a 3D grid search to optimize the cross-validated loss^87^. We masked 20% of the entries as held-out data, chosen in randomly selected 7-frame blocks (∼5.3 s) of consecutive time points in random neurons and trials. Blocked masking of held-out data was done to avoid temporal correlations between the training and testing data due to the slow dynamics of the calcium indicator. To protect further against spuriously high cross-validation performance due to temporal correlations, we trimmed the first 2 frames (∼1.5 s) from each block. These data were discarded from the test set, and only the remaining interior of each block was used to calculate the cross-validated loss. We repeated the grid search ten times with different random seeds for the train-test split and parameter initialization. We used a learning rate of 0.001, maximum iterations of 10000, and a minimum standard deviation of 10^-4^. We ran a non-constrained sliceTCA that allowed both positive and negative weights. Admissible models were defined as those achieving a minimum of 95% of the optimal performance. We selected the model by identifying the number of components that (1) generated a plateaued average test loss, (2) reconstructed all neurons with positive goodness-of-fit, and (3) had the highest sparsity across neuron weights.

#### Reconstruction performance

For each neuron, we estimated the goodness of fit of the sliceTCA reconstruction (**Figure 5G**)^87^:

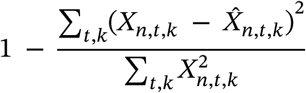

where 𝑋 is the input data and 𝑋^ is the reconstructed data at neuron 𝑛, time 𝑡, and trial 𝑘. This goodness of fit asks how much of the neuron’s total activity is explained by the reconstruction. However, if a neuron has high overall activity, it can achieve high goodness of fit even if the model just tracks mean trends. If a neuron exhibits low temporal variance, small reconstruction errors appear disproportionately large. Therefore, for model selection (**Figure S13C**), we compared the model error to the variance of each neuron’s activity:

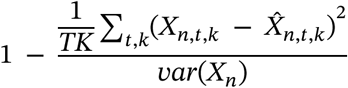

where T is the total number of time points and K is the total number of trials. With this, we instead asked how well the reconstruction explains fluctuations in activity compared to just using the average response. If this goodness of fit is negative, it indicates that the model is worse than predicting the mean. Therefore, models with negative values in this goodness of fit calculation are eliminated.

#### Sparsity

For model selection, we calculated the sparsity of the neuron component weights for each neuron (**Figure S13D**):

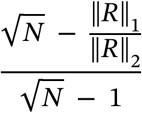

where 𝑁 is the number of neuron components, 𝑅 is the vector of component weights for a single neuron, is the sum of absolute values (L^1^ norm or Manhattan distance), and 𝑅 is the square root of the sum of squares (L^2^ norm or Euclidean distance):

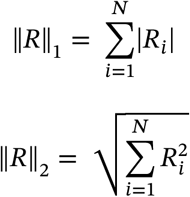

This sparsity metric ranges from 0 (dense) to 1 (sparse), with high sparsity when a neuron’s component weights are dominated by just a few nonzero entries.

#### Neuron clustering

Each neuron was assigned to the component for which it had the highest absolute neuron weight. Because weights can be positive or negative, each of the four components was divided into two clusters based on weight polarity, resulting in eight total clusters. To analyze temporal dynamics, neuronal responses were averaged within each cluster (**Figures 5J** and **S14**). Stimulus contributions to each cluster were assessed by plotting the distributions of absolute neuron weights using split violin plots (**Figure 5J**).

## Quantification and statistical analysis

Statistical analyses were conducted with Python scripts or GraphPad Prism (version 10.4.1). Detailed outline of our statistical analysis strategy can be found in **Table S3**. Data were evaluated for normality using the appropriate normality test. Normally distributed datasets were run with parametric statistical tests, whereas non-normally distributed datasets were run with nonparametric statistical tests. For comparing two datasets, we checked for the similarity of variances. For comparing more than two normally distributed datasets, we checked for the similarity of standard deviations using the Brown-Forsythe test.

For non-normal datasets with two independent variables without repeated measures, we fit a generalized linear model (GLM; **Table S2**). For comparing percentages (**Figures 1H**, **S10C**, and **S11C**), since they have a lower and upper bound (0-100%), we ran GLMs with a beta regression. The percentages were brought to be in the range (0, 1), and to avoid having exactly 0 and 1 values, we further transformed the data according to Smithson and Verkuilen (2006)^156^:

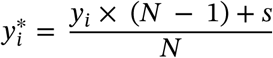

where 𝑦_𝑖_ ∈ [0, 1] is the original percentage value divided by 100, 𝑁 is the total number of values, 𝑠 ∈ (0, 1) is a small constant set to 0.5 for our calculations, and 𝑦^∗^ ∈ (0, 1) is the resulting value that avoids exact zeros or ones that the beta distribution cannot handle. For all other distributions, which are skewed and continuous, we used a Gamma GLM with a log link. Since some values, such as fold change (**Figures 1F** **and 1I**) or peak dF/F (**Figures S10E** and **S11G**) can be negative, all the values were shifted by the same number to be positive. A significant interaction or main effect was followed up with the appropriate multiple comparisons test. In our feeding assay, relative gut fluorescence values of WT and EEC^DTA^ had no significant differences between genotypes across stimuli, so we ran separate Kruskal-Wallis tests (**Figures 1F** **and 1I**; **Table S2**).

Results of statistical tests can be found on https://gitlab.oit.duke.edu/ean26/arinel-et-al_2025 and **Table S2**. All results are reported as mean ± standard error of the mean unless specified otherwise. Sample sizes were not predetermined by power analyses but were informed by preliminary experiments that evaluated the variance and effect sizes. Randomization and blinding to experimental conditions were not implemented during data collection or analysis.

