## Supplementary Figures for "Gut distension evokes rapid neural dynamics in vagal and hindbrain populations of larval zebrafish"

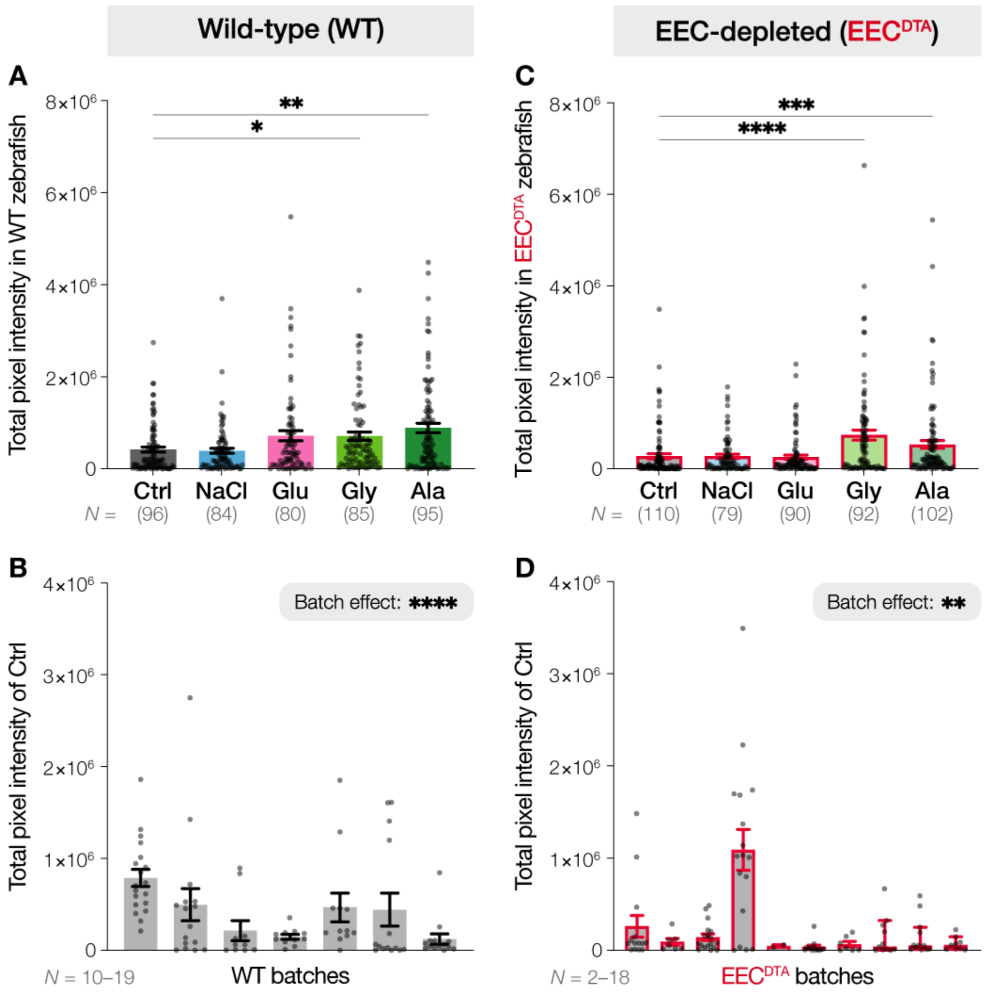


**Figure S1 | Batch-to-batch variability in feeding in larval zebrafish**

(**A**) Raw, total gut fluorescence in wild-type (WT) zebrafish fed with different complex particles (CPs). Comparing each CP with control (Ctrl, *N* = 96) showed higher feeding with glycine (Gly, *N* = 85, **P* = 0.0137) and alanine (Ala, *N* = 95, ***P* = 0.0031) in the gut, and no significant differences with NaCl (*N* = 84, *P* > 0.9999) or glucose (Glu, *N* = 80, *P* = 0.1595).
(**B**) Raw, total gut fluorescence to Ctrl particles across WT zebrafish batches (*N* = 10–19 fish per batch) had a significant batch effect (*****P* < 0.0001).
(**C**) Raw, total gut fluorescence in EEC-depleted ($EEC^{DTA}$) zebrafish fed with different CPs. Comparing each CP with Ctrl (*N* = 110) showed higher feeding with glycine (*N* = 92, *****P* < 0.0001) and alanine (*N* = 102, ****P* = 0.0004) in the gut, and no significant differences with NaCl (*N* = 79, *P* > 0.9999) or glucose (*N* = 90, *P* > 0.9999).
(**D**) Raw, total gut fluorescence to Ctrl particles across $EEC^{DTA}$ zebrafish batches (*N* = 2–18 fish per batch) had a significant batch effect (***P* = 0.0053).
Kruskal-Wallis and Dunn's multiple comparisons tests were run for statistical comparisons. Dots represent individual fish. Mean ± SEM.

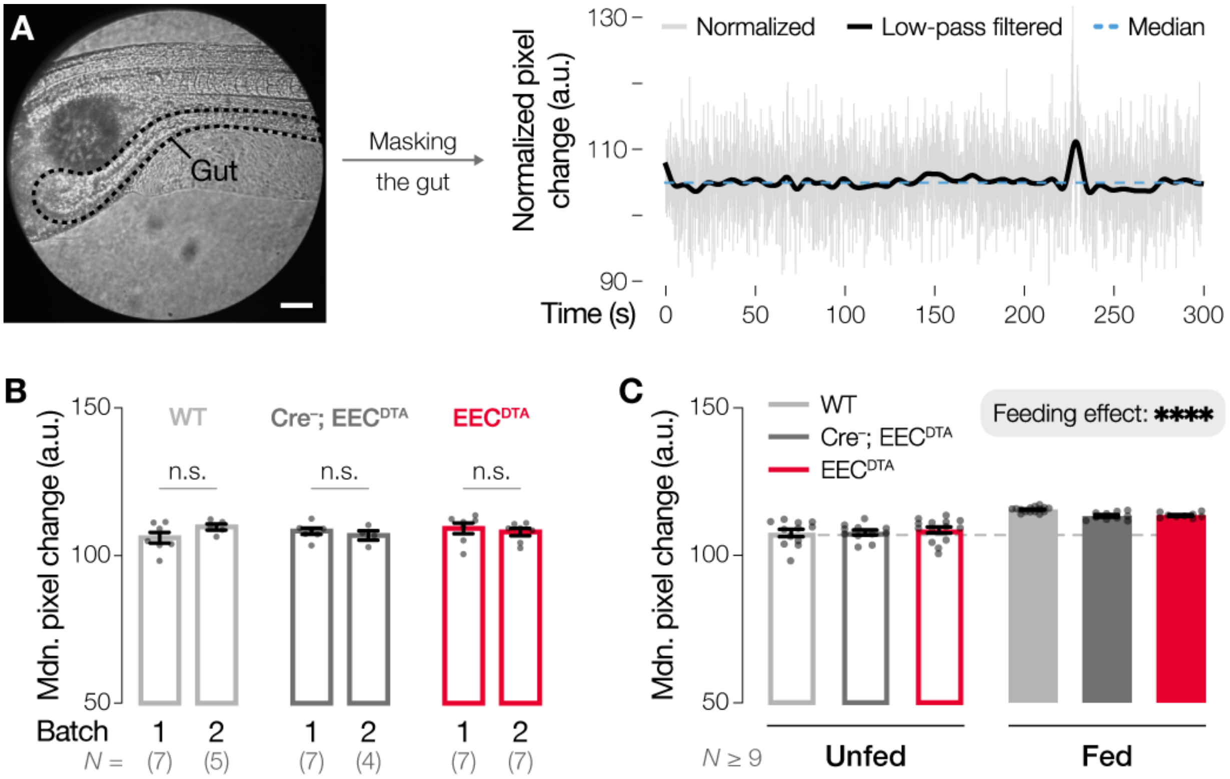


**Figure S2 | EEC$^{DTA}$ fish exhibit normal overall gut motility**

(**A**) Left: representative still frame from a gut motility recording of a 9 dpf, unfed, wild-type (WT) zebrafish, corresponding to Video S1. Dashed lines outline the gut, which was manually traced and masked for further analyses. Scale bar indicates 100 µm. Right: normalized pixel changes over time, which was calculated as the total pixel change within the gut mask divided by the gut area (light gray trace). To parse out gut motility, a low-pass Butterworth filter was applied (black trace). Due to twitch events (peak at ~230 seconds), we used the median of the filtered trace (blue dashed line) as a measure of gut motility.

(**B**) Median pixel changes of unfed fish from two independent experimental batches were compared across three genotypes: WT, Cre$^{-}$; EEC$^{DTA}$, EEC$^{DTA}$. There were no significant batch differences across any genotype ($N$ = 4–7, WT: $P$ = 0.3000, Cre$^{-}$; EEC$^{DTA}$: $P$ = 0.8562, EEC$^{DTA}$: $P$ = 0.9188, multiple unpaired t-tests with Welch and Šídák-Bonferroni corrections).

(**C**) Median pixel changes were compared between unfed (hollow bars) and fed (filled bars) fish across the three genotypes. The dashed gray line marks the average of unfed WT fish for reference. There were no significant differences across genotypes ($N$ = 9–15, $P$ = 0.5061), whereas feeding significantly increased gut motility (****$P$ < 0.0001, ordinary two-way ANOVA).

Dots represent individual fish (B and C). n.s., not significant. Mean ± SEM.

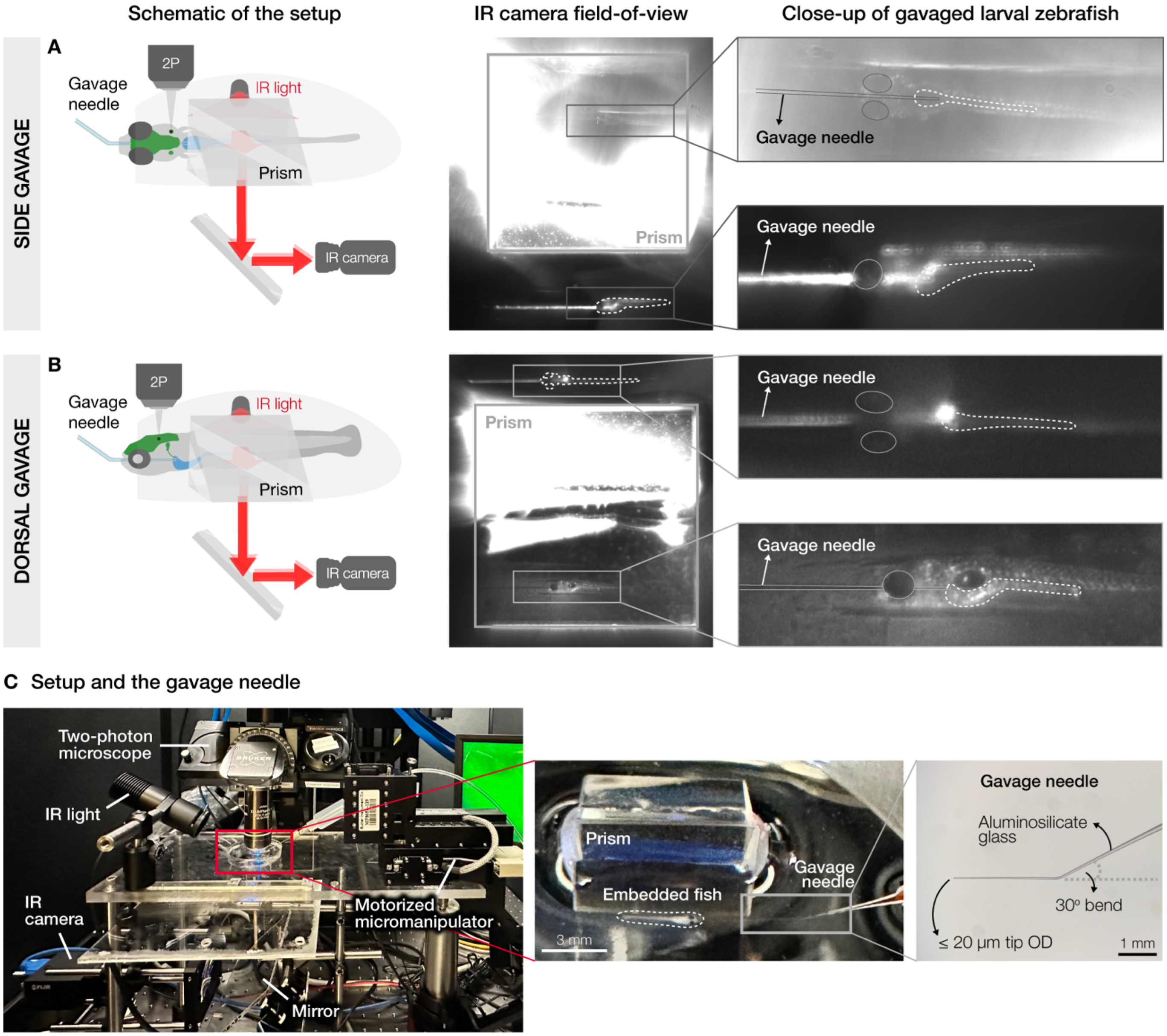


#### Figure S3 | Microgavage for enteric stimulus delivery during two-photon calcium imaging

(**A**) Left: schematic of the side gavage setup. A fish is embedded and gavaged sideways for vagal imaging. Middle: snapshot of the full field-of-view from the infrared (IR) camera. Prism mirror allows us to visualize all axes of the fish in a single frame. Right: zoom-in images of dorsal and side views for precise positioning of the needle. The gut is outlined (dashed lines).
(**B**) Left: schematic of the dorsal gavage setup. A fish is embedded and gavaged dorsal-side up for hindbrain imaging. Middle: snapshot of the full field-of-view from the IR camera. Right: zoom-in images of ventral and side views for precise positioning of the needle. The gut is outlined (dashed lines).
(**C**) Left: image of the experimental setup. A motorized micromanipulator enables precise positioning and controlled movement of the gavage needle. IR light directed at the fish under the objective is reflected by a mirror beneath the stage and captured by an IR camera. Although a FLIR camera is shown, the setup used for microgavage experiments employed a higher-resolution camera (1800 U-2050m, Allied Vision). Middle: closeup view of a gavaged larval zebrafish. A prism mirror positioned beside the fish reflects the lateral view to the IR camera. Right: closeup of the bent microgavage needle. Microgavage needles are prepared from aluminosilicate glass with a ≤20 µm outer diameter (OD) at the tip and a 30° bend 3–5 mm from the tip.

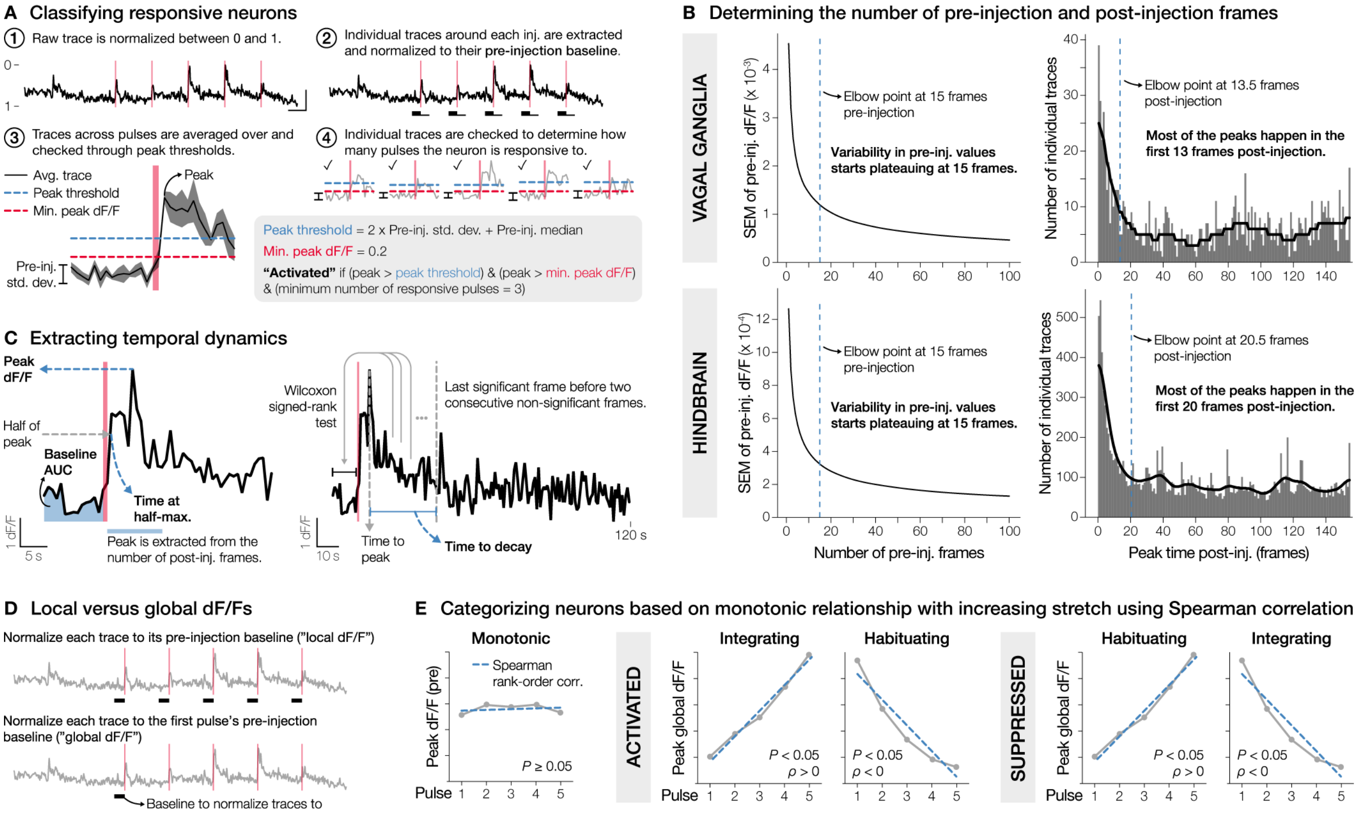


#### Figure S4 | Temporal analysis of two-photon calcium imaging data

(**A**) The analysis pipeline to identify responsive neurons consists of four steps. (1) Each neuron's raw fluorescence was normalized between 0 and 1. Scale bar: 0.5 dF/F on y-axis and 1 minute on x-axis. (2) Individual traces around each pulse were extracted from pre-injection (thick lines) to post-injection (thin lines) frames. Traces were normalized as dF/F to the median of pre-injection ("local dF/F"). (3) dF/F traces were averaged, and standard deviation of its pre-injection period was calculated. Peak threshold was defined as the pre-injection median plus two times its standard deviation. Since neurons with very stable baselines could be falsely classified as responsive due to minimal fluctuations, we implemented an additional minimum peak dF/F threshold of 0.2. To be classified as "activated", a neuron's post-injection peak had to exceed both the peak threshold and the minimum peak dF/F. Graph shows mean ± SEM. (4) If a neuron's average trace passed these thresholds, same analyses were repeated on individual pulse traces. A "responsive" neuron had at least 3 "responsive" traces.
(**B**) Left: determining the number of pre-injection frames. SEMs of pre-injection dF/F values across increasing frame lengths were calculated for vagal (top) and hindbrain (bottom) neurons. Number of pre-injection frames was set to the elbow point (dashed blue lines) of the histogram. Right: determining the number of post-injection frames. Distributions of peak times of all vagal (top) or hindbrain (bottom) neurons were calculated and smoothed with a Gaussian filter (solid black lines). Number of post-injection frames was set to the elbow point of the histogram.
(**C**) Left: representative dF/F trace, illustrating different temporal measures. Peak dF/F: maximum fluorescence in the post-injection window (blue bar). Time at half-maximum: time to reach half of peak dF/F, which was calculated by interpolating where the trace crossed the half-maximum value. Baseline AUC: baseline activity, calculated area under the curve (AUC) of the pre-injection period. Right: same response trace plotted over 120 seconds to determine time to decay, which was defined as the duration from peak until fluorescence returned to baseline. Each post-peak frame was compared to the pre-injection period with Wilcoxon signed-rank test. If two consecutive frames were not significantly different from baseline, the time from peak to the last significant frame was recorded as time to decay.
(**D**) Two different baselines (black bars) were used for normalization of individual traces. Top: "local dF/F" normalizes to the median fluorescence of each trace's own pre-injection period. Bottom: "global dF/F" normalizes

to the median fluorescence of first pulse's pre-injection period. Local dF/Fs were used for time at half-maximum and time to decay calculations, while global dF/Fs were used for baseline AUC measurements and neuron classifications by responses to increasing distension. Peak dF/F values were plotted using either normalization, as specified in individual figures.

(**E**) To determine whether neurons encoded stretch intensity, we extracted peak global dF/F values and ran a Spearman rank-order correlation to test for a significant relationship ($P < 0.05$). Neurons with no significant correlation were classified as monotonic responders. Activated neurons with a significant correlation were labeled as integrating if their responses increased over pulses ($\rho > 0$) or habituating if their responses decreased ($\rho < 0$), and vice versa for suppressed neurons. $\rho$ indicates the Spearman correlation coefficient.

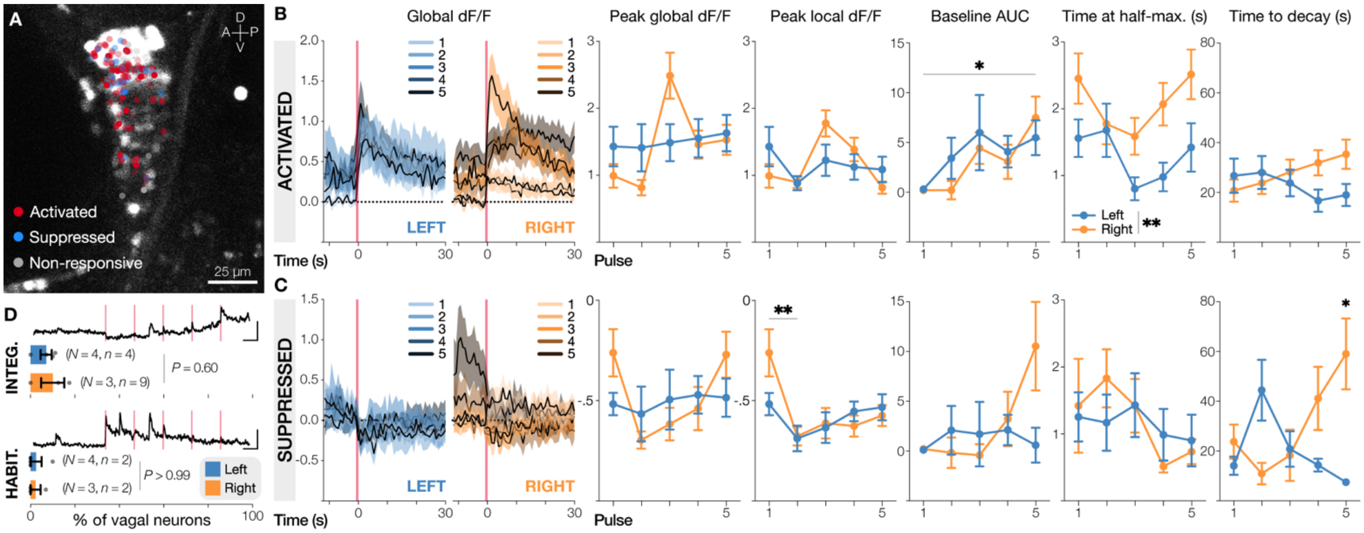

**Figure S5 | Vagal neuron dynamics to increasing distension exhibit minimal lateralization**

(**A**) Spatial distribution of vagal neurons classified by response direction. Neurons from all five imaging planes were overlaid onto a left vagal ganglion reference image. Neurons from the right vagus were horizontally flipped for overlay. Because cross-fish alignment could not reliably distinguish posterior lateral line ganglion from vagal ganglion, all neurons within the ganglionic region were included for analysis. Dot opaqueness indicates higher average peak local dF/F across pulses. A: anterior, D: dorsal, P: posterior, V: ventral.
(**B**) Temporal dynamics of activated neurons (left vagus: blue, *n* = 31 neurons, right vagus: orange, *n* = 55 neurons). Left: average dF/F traces across five pulses, normalized to the baseline of the first pulse ("global dF/F"). Darker traces represent increasing distension. Right: peak global dF/Fs, peak local dF/Fs, baseline AUCs, times at half-maximum, and times to decay of activated neurons across pulses. Time at half-maximum: right vagal neurons had slower onsets than left (***P* = 0.0056, mixed-effects analysis).
(**C**) Temporal dynamics of suppressed neurons (left vagus: *n* = 13 neurons, right vagus: *n* = 12 neurons), similar to (B). Time to decay: right vagal neurons were suppressed longer at higher distension intensities compared to left (**P* = 0.0207, mixed-effects analysis and Šídák's multiple comparisons test).
(**D**) Representative normalized dF/F traces and percentages of integrating (top) or habituating (bottom) vagal neurons, which were similar between left and right vagus (integrating: unpaired t-test, habituating: Mann-Whitney test). Scale bar for traces: 0.5 dF/F on y-axis and 1 minute on x-axis. Dots represent individual fish (*N*), with the total neuron count (*n*) shown.
Injection times are marked by red lines. Mean ± SEM.

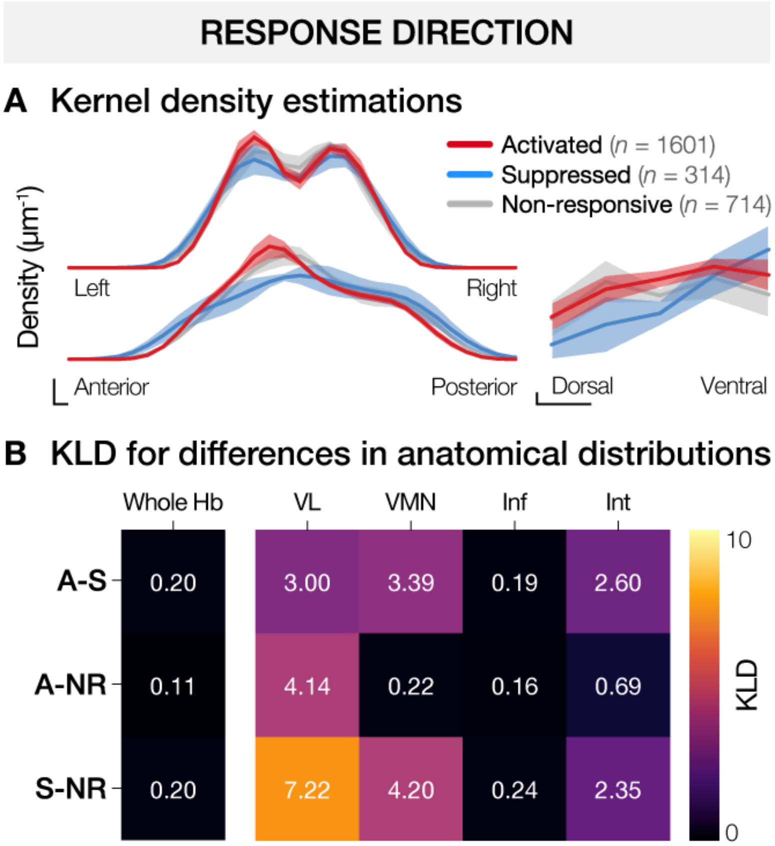


**Figure S6 | Stretch-responsive hindbrain neurons do not show spatial organization**

Spatial distributions of neurons classified by response type: activated (A, red, *n* = 1601 neurons, *N* = 4 fish), suppressed (S, blue, *n* = 314 neurons), and non-responsive (NR, gray, *n* = 714 neurons).
(**A**) Kernel density estimations illustrate neuron distributions along the left-right (top left), anterior-posterior (bottom left), and dorsoventral (right) anatomical axes. Comparisons of the area under the kernel density estimation curves revealed no significant differences between left (0–250 μm) vs. right (250–500 μm, *P* = 0.4680, repeated measures two-way ANOVA), anterior (0–250 μm) vs. posterior (250–500 μm, *P* = 0.1410, repeated measures two-way ANOVA), and dorsal (0–20 μm) vs. ventral (20–40 μm, *P* = 0.2371, mixed-effects analysis) distributions. Scale bars: 0.002 $\mu m^{-1}$ (y-axis), 10 μm (x-axis).
(**B**) Heatmap depicting Kullback-Leibler divergences (KLD) quantify spatial divergence between pairwise comparisons of neuron response types across the hindbrain (left) and within specific brain regions (right). Higher KLD values indicate greater differences in anatomical distributions. Left: across the entire hindbrain, KLD values showed a trending separation across neuron types (*P* = 0.0696, repeated measures one-way ANOVA). Right: within specific regions, there were no significant differences in KLDs by region (*P* = 0.2547) or response direction (*P* = 0.2960, mixed-effects analysis). A: activated, S: suppressed, NR: non-responsive, Hb: hindbrain, VL: vagal sensory lobe, VMN: vagus motor nucleus, Inf: inferior dorsal medulla oblongata, Int: intermediate dorsal medulla oblongata.
Mean ± SEM.

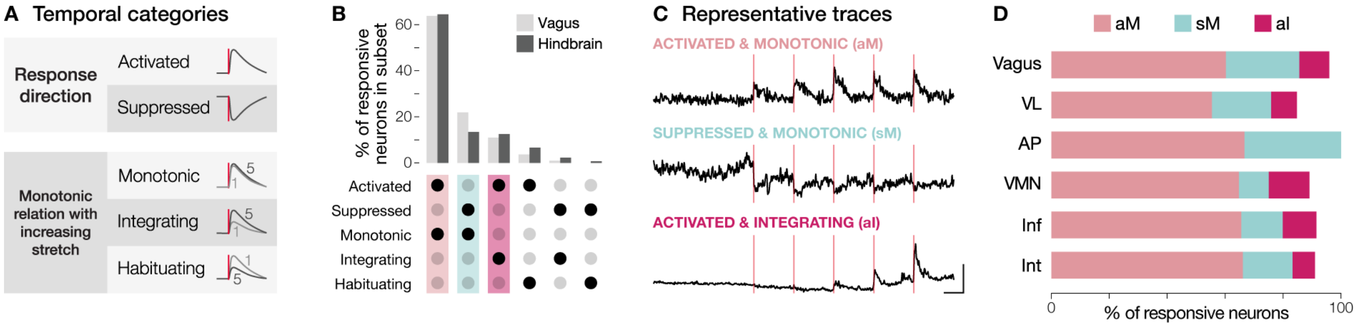


#### Figure S7 | Temporal classification of distension-responsive vagal and hindbrain neurons

(**A**) Temporal categories based on response direction and peak global dF/F relationship with increasing distension (see Figure S4E).
(**B**) UpSet plot showing the percentage of responsive vagal (light gray bars) and hindbrain (dark gray bars) neurons across temporal categories. Black circles indicate inclusion in a subtype. We selected vagal subtypes with at least 10 neurons in total and representation in at least half of the recorded fish, which yielded three prominent categories: activated & monotonic (aM, light pink), suppressed & monotonic (sM, light blue), and activated & integrating (a**I**, pink). For hindbrain, we selected subtypes with at least 10 neurons from each fish, which identified aM, sM, a**I**, and activated & habituating neurons. Given the overlap, aM, sM, and a**I** were chosen for further analyses.
(**C**) Representative normalized dF/F traces of hindbrain neurons from each subtype. Scale bar for traces: 0.5 dF/F on y-axis and 1 minute on x-axis. Injection times are marked by red lines.
(**D**) Average percentages of responsive neurons broken down by temporal subtype, which were similar across vagal and hindbrain regions (aM: $P$ = 0.9584, Brown-Forsythe test, sM: $P$ = 0.4113, Kruskal-Wallis test; a**I**: $P$ = 0.1438, Kruskal-Wallis test). AP: area postrema, Inf: inferior dorsal medulla oblongata, Int: intermediate dorsal medulla oblongata, VL: vagal sensory lobe, VMN: vagus motor nucleus.Bars show mean.

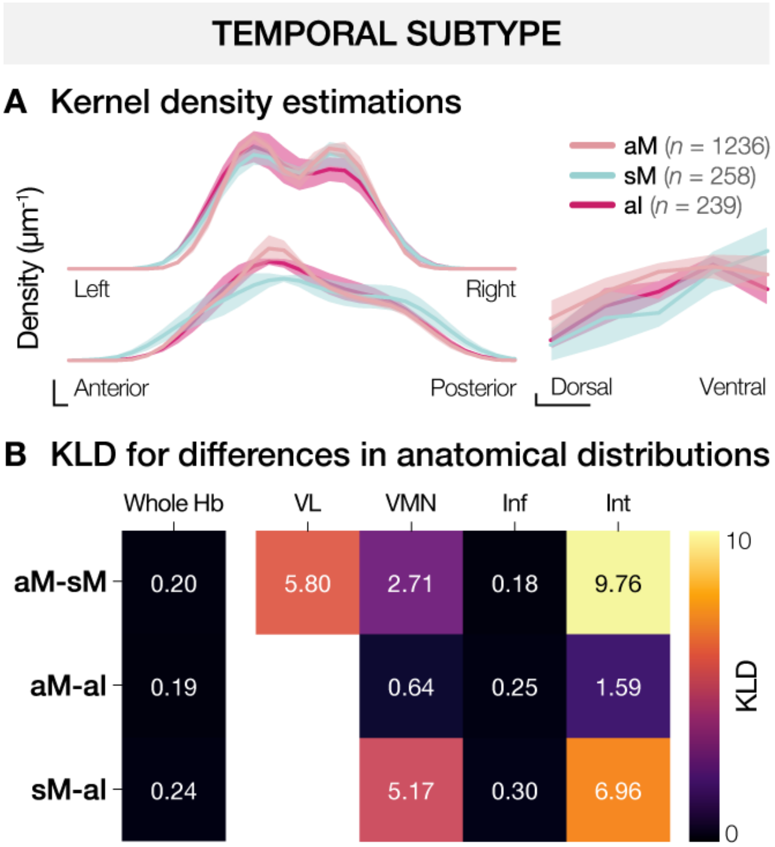


**Figure S8 | Temporal subtypes are broadly distributed across the dorsal hindbrain**

Spatial distribution of activated & monotonic (aM, light pink, *n* = 1236 neurons, *N* = 4 fish), suppressed & monotonic (sM, light blue, *n* = 258 neurons), and activated & integrating (aI, pink, *n* = 239 neurons) neuronal subtypes.
(**A**) Kernel density estimations illustrate neuron distributions along the left-right (top left), anterior-posterior (bottom left), and dorsoventral (right) anatomical axes. Comparisons of kernel density estimation area under the curve indicated no significant differences between left vs. right (*P* = 0.8219, repeated measures two-way ANOVA), anterior vs. posterior (*P* = 0.4767, repeated measures two-way ANOVA), and dorsal vs. ventral (*P* = 0.0617, mixed-effects analysis) axes. Scale bars: 0.002 $\mu m^{-1}$ (y-axis), 10 µm (x-axis).
(**B**) Heatmap depicting Kullback-Leibler divergences (KLD) quantify spatial divergence between pairwise comparisons of temporal subtypes across the hindbrain (left) and within specific brain regions (right). Higher KLD values indicate greater differences in anatomical distributions. Left: across the entire hindbrain, there were no significant differences in KLDs among subtypes (*P* = 0.2731, Friedman test). Right: within individual brain regions, there were no significant differences in spatial divergence by region (*P* = 0.2479) or neuronal subtype (*P* = 0.3346, mixed-effects analysis). Due to insufficient numbers of aI neurons in VL, KLDs involving this subtype (aM-aI, sM-aI) could not be calculated and thus were omitted from the heatmap. Hb: hindbrain, VL: vagal sensory lobe, VMN: vagus motor nucleus, Inf: inferior dorsal medulla oblongata, Int: intermediate dorsal medulla oblongata. Mean ± SEM.

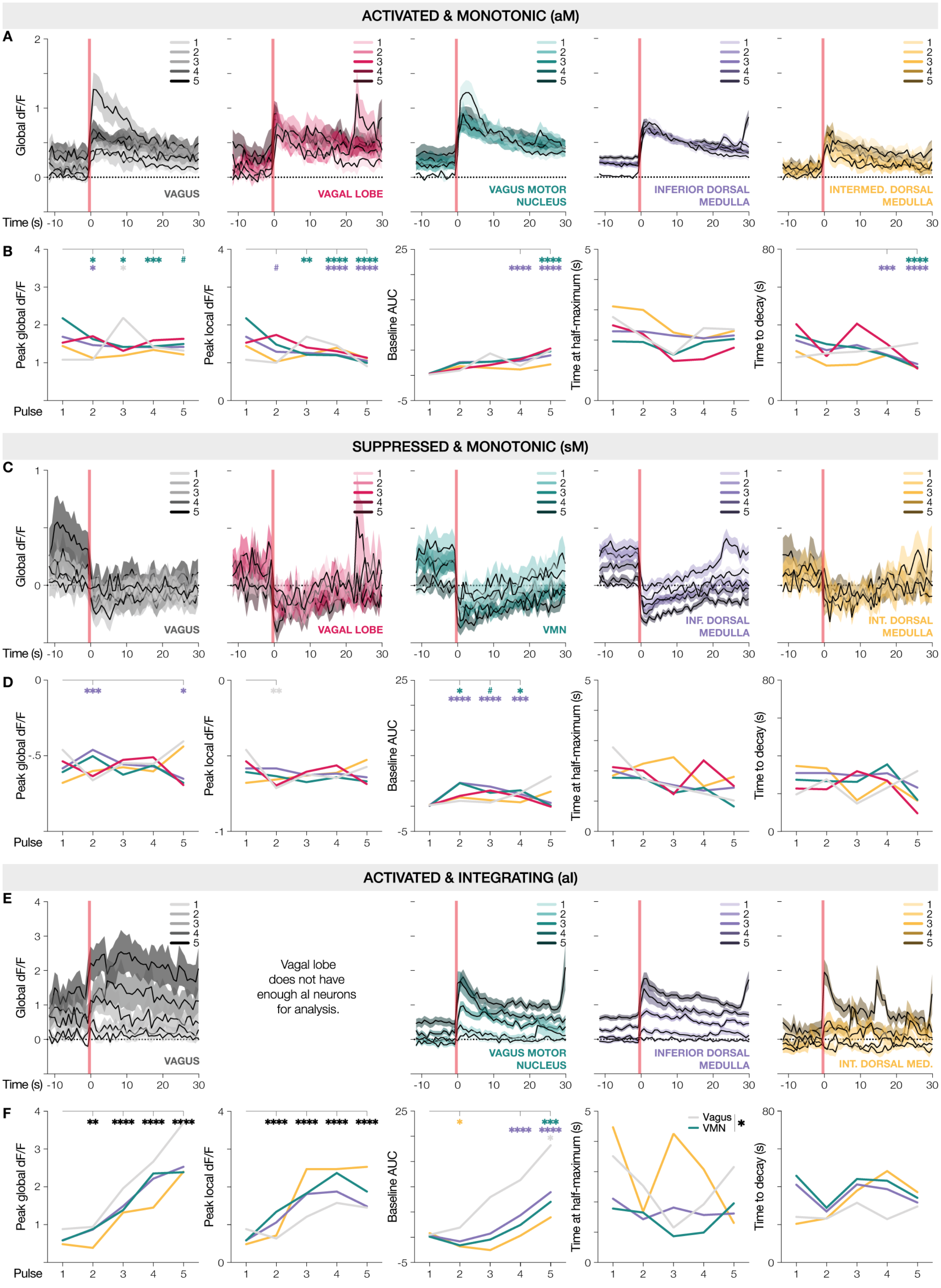

ACTIVATED & MONOTONIC (aM)
A
Global dF/F
Time (s)
VAGUS
VAGAL LOBE
VAGUS MOTOR NUCLEUS
INFERIOR DORSAL MEDULLA
INTERMED. DORSAL MEDULLA
B
Peak global dF/F
Peak local dF/F
Baseline AUC
Time at half-maximum (s)
Time to decay (s)
Pulse
SUPPRESSED & MONOTONIC (sM)
C
VMN
INF. DORSAL MEDULLA
INT. DORSAL MEDULLA
D
ACTIVATED & INTEGRATING (aI)
E
Vagal lobe does not have enough al neurons for analysis.
INT. DORSAL MED.
F
Vagus
VMN

#### Figure S9 | Subtype dynamics to increasing distension across vagal and hindbrain regions

(**A**) Average dF/F traces of activated & monotonic (aM) neurons across five pulses, normalized to the baseline of the first pulse ("global dF/F"; vagus: gray, $n$ = 70 neurons, vagal sensory lobe: VL, pink, $n$ = 43 neurons, vagus motor nucleus: VMN, green, $n$ = 216 neurons, inferior dorsal medulla oblongata: Inf, purple, $n$ = 692 neurons, and intermediate dorsal medulla oblongata: Int, yellow, $n$ = 77 neurons). Darker traces represent increasing distension.
(**B**) Peak global dF/Fs, peak local dF/Fs, baseline AUCs, times at half-maximum, and times to decay of aM neurons across pulses. Peak global dF/F: VMN neurons had smaller peaks with increasing distension ($P$ < 0.03, $^{\#}P$ = 0.0849). Peak local dF/F: VMN and Inf neurons had smaller changes in peak with increasing distension ($P$ < 0.006, $^{\#}P$ = 0.0843). Baseline AUC: VMN and Inf neurons had increasing baseline activity (****$P$ < 0.0001). Time to decay: VMN and Inf neurons had shorter responses with increasing distension ($P$ < 0.0004).
(**C**) Average dF/F traces of suppressed & monotonic (sM) neurons across five pulses (vagus: $n$ = 24 neurons, VL: $n$ = 14 neurons, VMN: $n$ = 38 neurons, Inf: $n$ = 155 neurons, and Int: $n$ = 21 neurons).
(**D**) Baseline AUC: VMN and Inf neurons had increased baseline activity after the first pulse ($P$ < 0.05, $^{\#}P$ = 0.0760).
(**E**) Average dF/F traces of activated & integrating (aI) neurons across five pulses (vagus: $n$ = 12 neurons, VMN: $n$ = 46 neurons, Inf: $n$ = 126 neurons, and Int: $n$ = 10 neurons). VL is excluded due to insufficient number of neurons ($n$ = 6 neurons).
(**F**) Peak global dF/F: all regions had increasing peaks with increasing distension ($P$ < 0.002, Friedman test). Peak local dF/F: all regions had increasing change in peak with increasing distension (****$P$ < 0.0001, Friedman test). Baseline AUC: vagal, VMN, and Inf neurons had increasing baseline activity ($P$ < 0.03). Time at half-maximum: VMN neurons had faster onsets than vagal neurons (*$P$ = 0.0225, Kruskal-Wallis test).
Mixed-effects analyses with Šídák's multiple comparisons tests were run for statistical comparisons unless stated otherwise. Injection times are marked by red lines (A, C, and E). *$P$ < 0.05, **$P$ < 0.01, ***$P$ < 0.001, ****$P$ < 0.0001. Mean ± SEM.

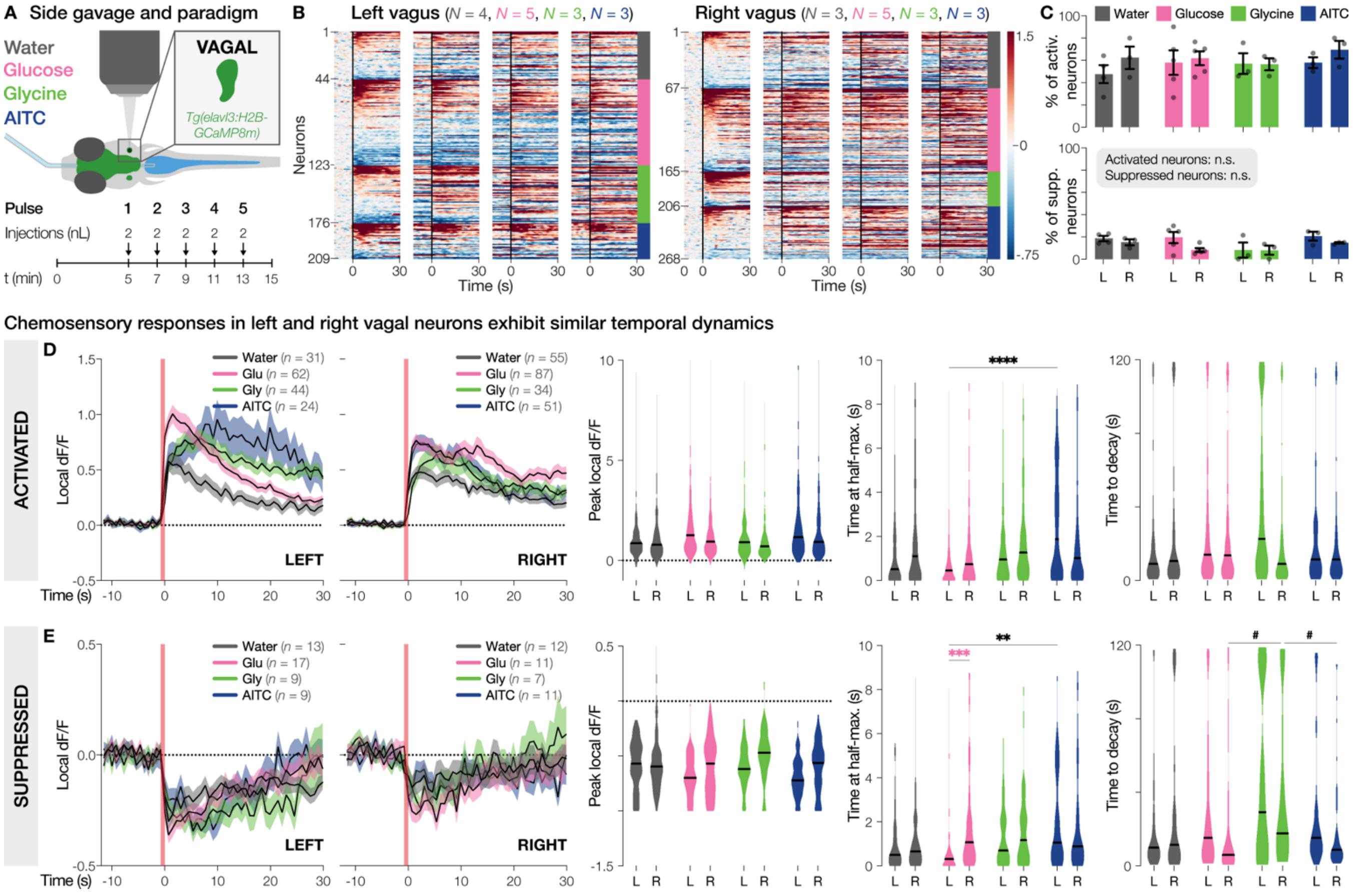


#### Figure S10 | Left and right vagal neurons exhibit similar encoding of enteric chemical stimuli

(**A**) Experimental setup. 6–10 dpf *Tg(elavl3:H2B-GCaMP8m)* zebrafish were embedded on their side and continuously imaged during microgavage with water, glucose (25 mM), glycine (25 mM), or AITC (25 mM), delivering five pulses of 2 nL.
(**B**) Heatmaps of all stimulus-responsive neurons from left (water: gray, $N = 4$ fish, glucose: pink, $N = 5$ fish, glycine: green, $N = 3$ fish, AITC: blue, $N = 3$ fish) and right (water: $N = 3$ fish; glucose: $N = 5$ fish; glycine: $N = 3$ fish; AITC: $N = 3$ fish) vagal ganglia across pulses, sorted by peak response to the first pulse. Black lines indicate injection pulses. Color scale indicates global dF/F.
(**C**) Percentages of activated (top) and suppressed (bottom) vagal neurons, which were similar between left and right vagus and across stimuli (activated: two-way ANOVA, suppressed: generalized linear model and Kruskal-Wallis test). Dots represent individual fish.
(**D**) Temporal dynamics of activated neurons around injections. Left: average dF/F traces from vagal neurons responsive to water, glucose (Glu), glycine (Gly), or AITC. Numbers in legends indicate number of neurons. Right: peak local dF/Fs, times at half-maximum, and times to decay of activated neurons. In the left vagus, AITC-responsive neurons had slower onsets than glucose-responsive ones (****$P < 0.0001$, $|\delta| = 0.3659$).
(**E**) Temporal dynamics of suppressed neurons. In the left vagus, AITC-responsive neurons had slower onsets than glucose-responsive ones (**$P = 0.0020$, $|\delta| = 0.4216$). Glucose-responsive right vagal neurons also had slower onsets than left (***$P = 0.0009$, $|\delta| = 0.3875$).
Mixed-effects analyses with Šídák's multiple comparisons tests and Cliff's delta were run for statistical comparisons unless stated otherwise. Injection times are marked by red lines (D and E). n.s., not significant. Line and bar graphs show mean ± SEM. Violin plots show median.

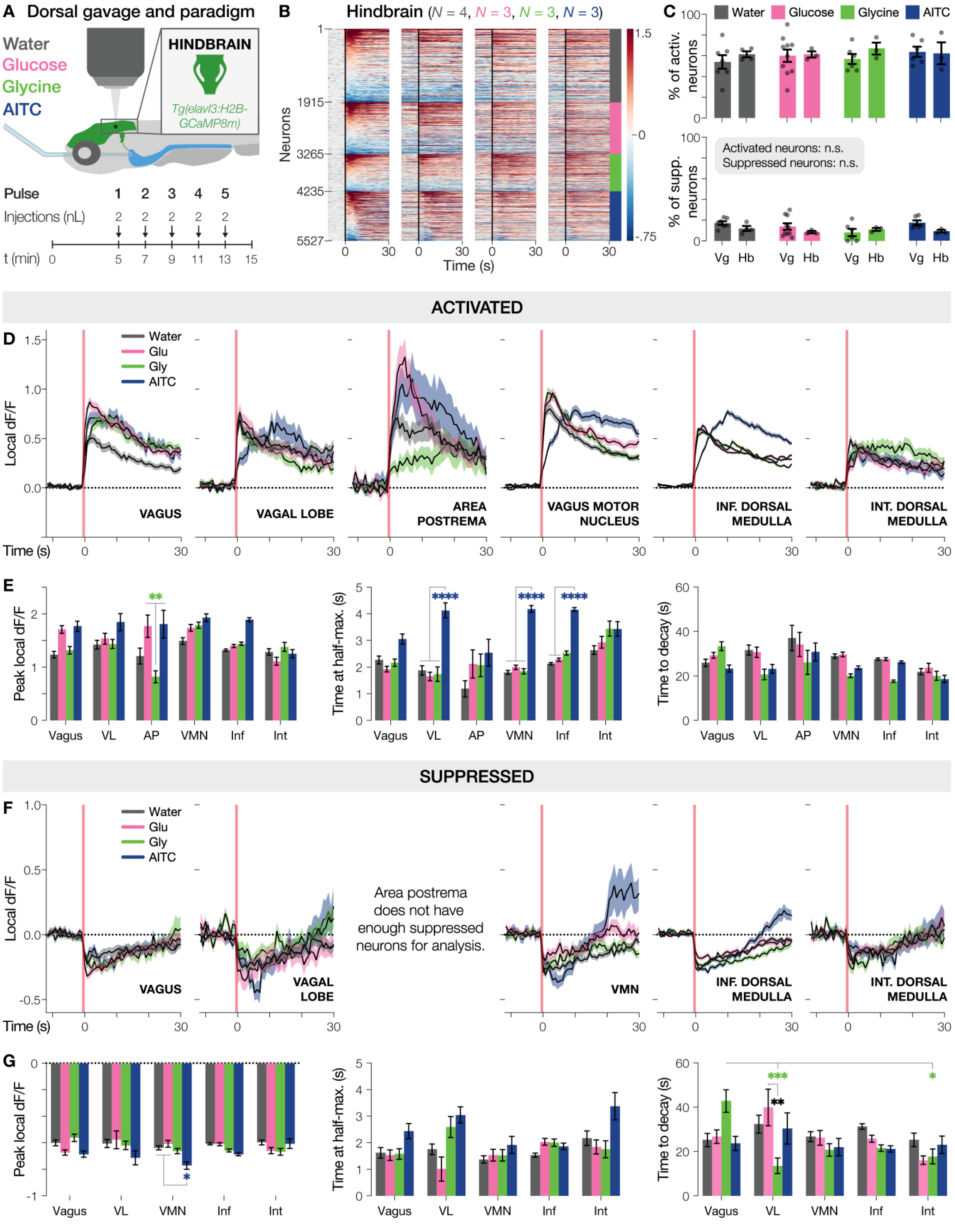

A Dorsal gavage and paradigm
Water
Glucose
Glycine
AITC
HINDBRAIN
Tg(elavl3:H2B-GCaMP8m)
Pulse 1 2 3 4 5
Injections (nL) 2 2 2 2 2
t (min) 0 5 7 9 11 13 15
B
Hindbrain (N = 4, N = 3, N = 3, N = 3)
Neurons
Time (s)
C
Water Glucose Glycine AITC
% of activ. neurons
% of supp. neurons
Activated neurons: n.s.
Suppressed neurons: n.s.
Vg Hb Vg Hb Vg Hb Vg Hb
ACTIVATED
D
Water
Glu
Gly
AITC
Local dF/F
VAGUS
VAGAL LOBE
AREA POSTREMA
VAGUS MOTOR NUCLEUS
INF. DORSAL MEDULLA
INT. DORSAL MEDULLA
Time (s)
E
Peak local dF/F
Time at half-max. (s)
Time to decay (s)
Vagus VL AP VMN Inf Int
SUPPRESSED
F
Water
Glu
Gly
AITC
Local dF/F
VAGUS
VAGAL LOBE
Area postrema does not have enough suppressed neurons for analysis.
VMN
INF. DORSAL MEDULLA
INT. DORSAL MEDULLA
Time (s)
G
Peak local dF/F
Time at half-max. (s)
Time to decay (s)
Vagus VL VMN Inf Int

#### Figure S11 | AITC evokes hindbrain responses with slower onsets

(**A**) Experimental setup. 6–10 dpf *Tg(elavl3:H2B-GCaMP8m)* zebrafish were embedded dorsal-side-up and continuously imaged during microgavage with water, glucose (25 mM), glycine (25 mM), or AITC (25 mM), delivering five pulses of 2 nL.
(**B**) Heatmap of all stimulus-responsive hindbrain neurons across pulses, sorted by peak response to the first pulse (water: gray, $N$ = 4 fish, glucose: pink, $N$ = 3 fish, glycine: $N$ = green, 3 fish, AITC: blue, $N$ = 3 fish). Black lines indicate injection pulses. Color scale indicates global dF/F.
(**C**) Percentages of activated (top) and suppressed (bottom) neurons, which were similar between vagus and hindbrain and across stimuli (activated: two-way ANOVA, suppressed: generalized linear model and Šídák's multiple comparisons test). Dots represent individual fish.
(**D**) Average dF/F traces of activated neurons responsive to each stimulus.
(**E**) Peak local dF/Fs, times at half-maximum, and times to decay of activated neurons. Time at half-maximum: AITC-responsive neurons had slower onsets than other stimulus-responsive neurons in vagal sensory lobe (VL), vagus motor nucleus (VMN), and inferior dorsal medulla oblongata (Inf, ****$P$ < 0.0001, |δ| > 0.3525).
(**F**) Average dF/F traces of suppressed neurons responsive to each stimulus. Area postrema (AP) is excluded due to insufficient number of neurons (water: $n$ = 3 neurons, glucose: $n$ = 0 neurons, glycine: $n$ = 1 neuron, AITC: $n$ = 2 neurons).
(**G**) Peak local dF/Fs, times at half-maximum, and times to decay of suppressed neurons. Peak local dF/F: AITC-responsive vagus motor nucleus (VMN) neurons had larger peaks than water- ($P$ = 0.0057, |δ| = 0.3549) and glucose-responsive neurons (*$P$ = 0.0166, |δ| = 0.3535). Time to decay: glycine-responsive VL neurons were suppressed shorter than glycine-responsive vagal (***$P$ = 0.0009, |δ| = 0.4886) and glucose-responsive VL neurons (**$P$ = 0.0079, |δ| = 0.5513).
Mixed-effects analyses with Šídák's multiple comparisons tests and Cliff's delta were run for statistical comparisons unless stated otherwise. Injection times are marked by red lines (D and F). n.s., not significant. Mean ± SEM.

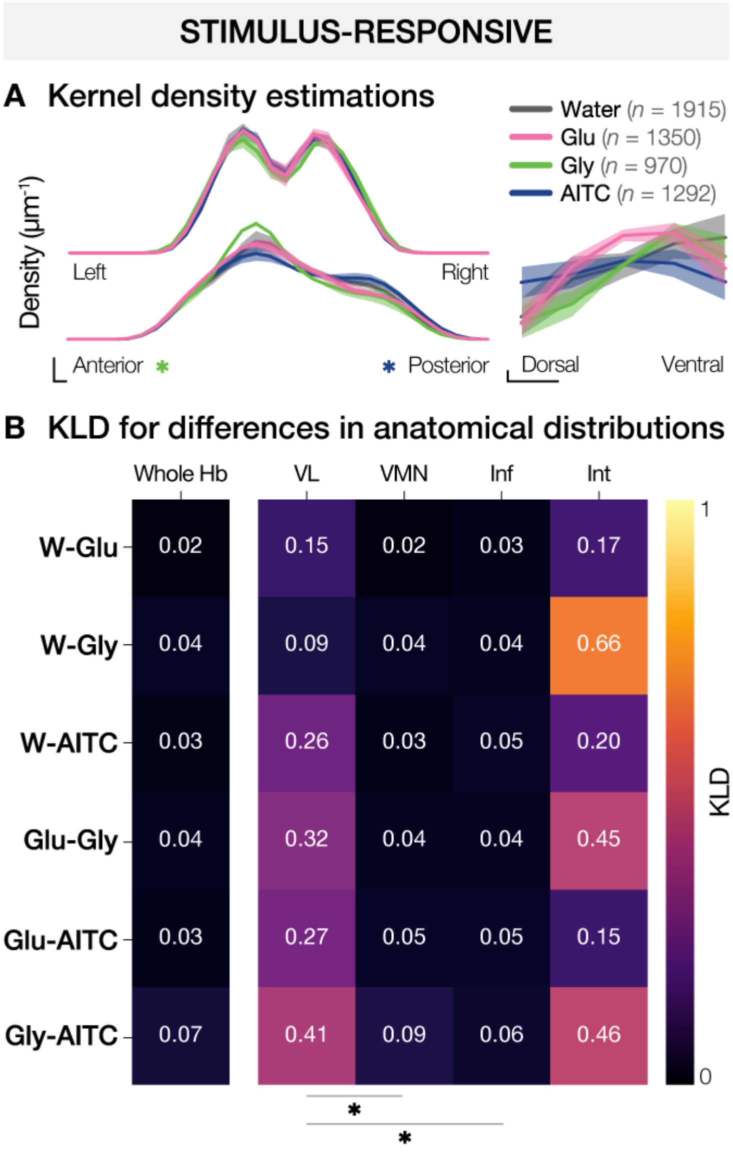


**Figure S12 | Glycine- and AITC-responsive neurons show opposing anterior-posterior biases**

Spatial distribution of neurons responsive to water (gray, $n$ = 1915 neurons, $N$ = 4 fish), glucose (Glu, pink, $n$ = 1350 neurons, $N$ = 3 fish), glycine (Gly, green, $n$ = 970 neurons, $N$ = 3 fish), and AITC (blue, $n$ = 1292 neurons, $N$ = 3 fish).
(**A**) Kernel density estimations illustrate neuron distributions along the left-right (top left), anterior-posterior (bottom left), and dorsoventral (right) anatomical axes. Along the anterior-posterior axis, glycine-responsive neurons were localized more anteriorly (*$P$ = 0.0458), while AITC-responsive neurons were shifted posteriorly (*$P$ = 0.0102, mixed-effects analysis and Šídák's multiple comparisons test). Scale bars: 0.002 µm$^{-1}$ (y-axis), 10 µm (x-axis).
(**B**) Heatmap depicting Kullback-Leibler divergences (KLD) quantify spatial divergence between pairwise comparisons of stimulus-responsive neurons across the hindbrain (Hb, left) and within specific brain regions (right). Higher KLD values indicate greater differences in anatomical distributions. Vagal sensory lobe (VL) had significantly higher divergence across stimuli compared to the vagus motor nucleus (VMN, *$P$ = 0.0263) and inferior dorsal medulla oblongata (Inf, *$P$ = 0.0352). Within VL, the KLD between glycine and AITC showed a trend toward greater divergence than other pairwise comparisons ($P$ = 0.0625, Wilcoxon signed-rank test). Inf: inferior dorsal medulla oblongata, Int: intermediate dorsal medulla oblongata.
Mean ± SEM.

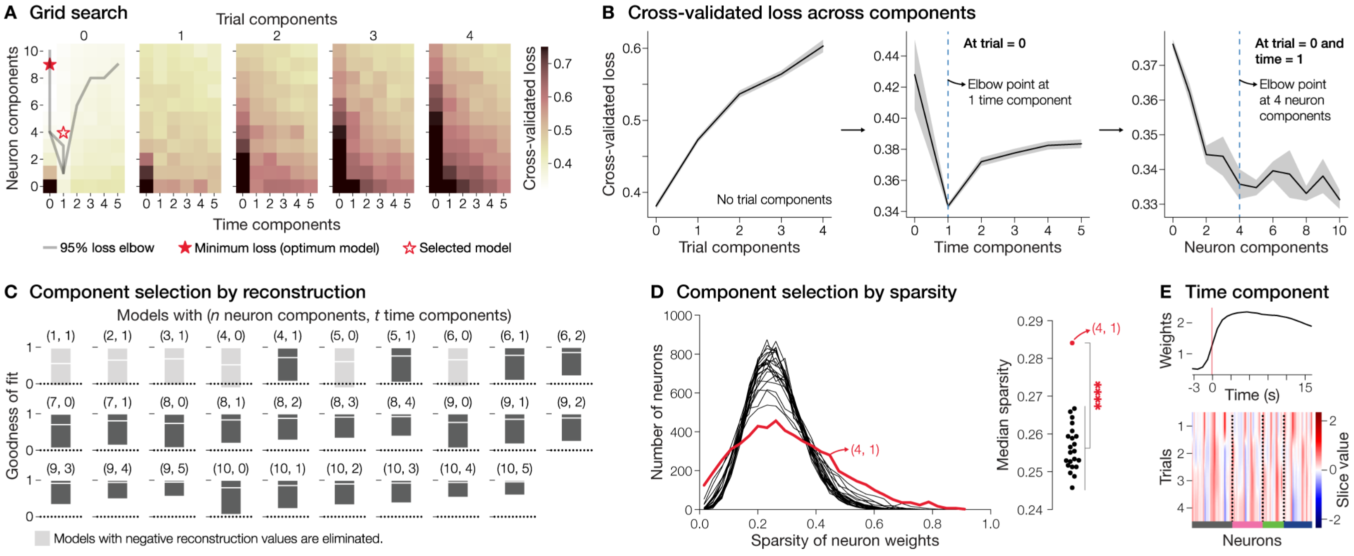


### Figure S13 | SliceTCA component selection

(**A**) Grid search to select optimal trial-, neuron-, and time-slicing components for the slice tensor component analysis (sliceTCA) model. Cross-validated loss was computed for combinations of trial, neuron, and time components. Optimal model with the lowest loss (filled star) and the selected model in Fig. 5 (hollow star) are shown. Gray lines represent the 95% loss elbow.
(**B**) Average cross-validated loss as a function of trial (left), time (middle), and neuron (right) components. Adding trial-slicing components increased overfitting, therefore, no trial components were included. For models without trial components, the elbow points of the losses were 1 time component and 4 neuron components. Graphs show mean ± SEM.
(**C**) Reconstruction performances (see Methods) of models with losses at the 95% elbow or lower. Models with negative reconstruction scores (light gray) were excluded from future analyses. Graphs show median ± range.
(**D**) Sparsity of neuron weights across components was compared among remaining models. Higher sparsity indicates more separable neuron clustering. Left: sparsity histograms of each model, with the most right-shifted model (4 neuron, 1 time component) in red. Right: median sparsity values, highlighting significantly greater sparsity in the (4, 1) model compared to other models (*****P* < 0.0001, Wilcoxon signed-rank test). Therefore, we continued with this 0-trial, 4-neuron, and 1-time component model.
(**E**) Weights (top) and slice (bottom) for the time-slicing component of the final model. As expected, the weight profile showed increased calcium activity post-injection, aligning with gut stimulus delivery (red line). Neurons in the slice heatmap are organized by stimulus responsivity (dotted black lines, water: gray, glucose: pink, glycine: green, AITC: blue).

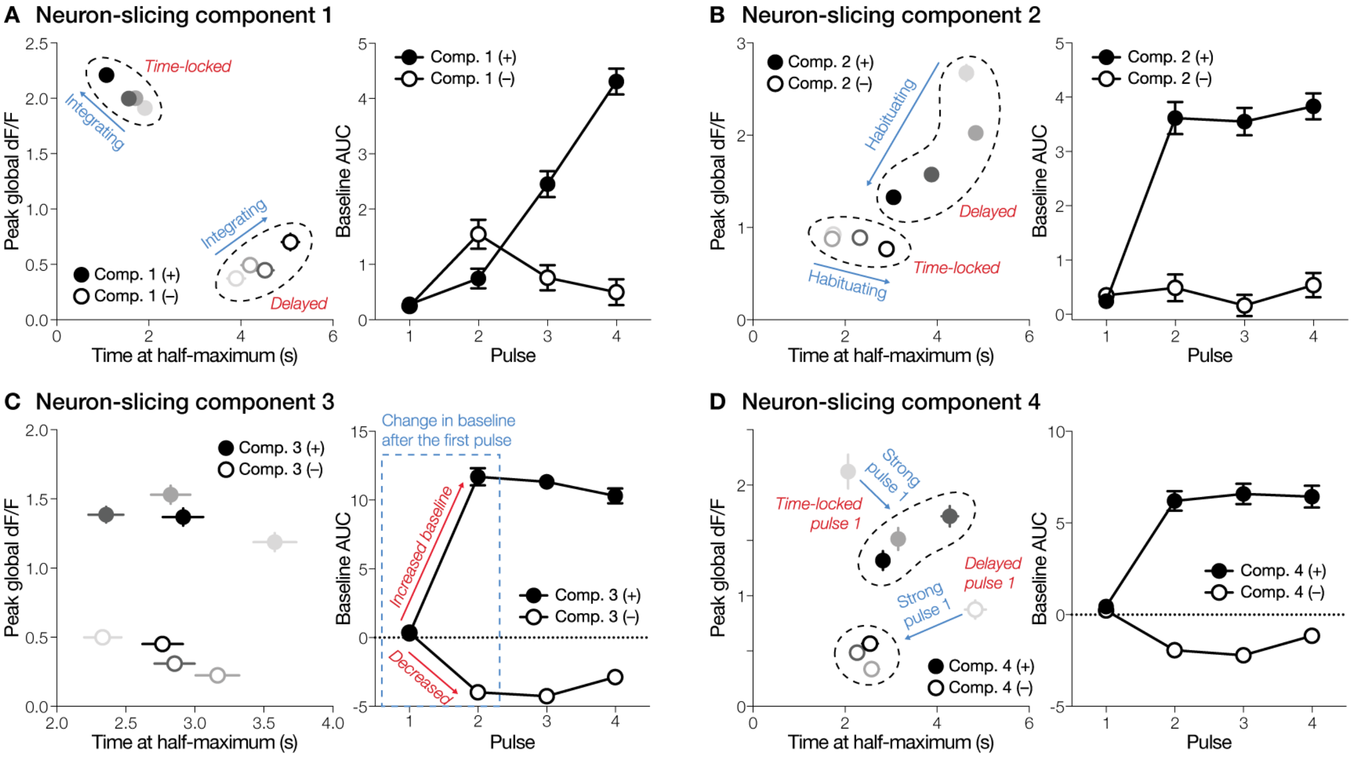


#### Figure S14 | Temporal identifiers of neuron-slicing components

For each neuron-slicing component, average peak global dF/F against time at half-maximum (left) and baseline area under the curve (AUC) against pulses (right) were plotted, to highlight common (blue) and distinct (red) temporal features between positively (filled circles) and negatively (open circles) weighted neuron clusters. Darker data points represent increasing pulses.

(**A**) Component 1 captured integrating neurons with increasing peak responses across pulses. Positive and negative weights represented time-locked versus delayed neurons, respectively.

(**B**) Component 2 captured habituating neurons with decreasing peak responses across pulses. Positive and negative weights represented delayed versus time-locked neurons, respectively.

(**C**) Component 3 captured neurons exhibiting pronounced baseline changes after the first pulse. Positive and negative weights corresponded to increased versus decreased baseline activity, respectively.

(**D**) Component 4 captured neurons with reduced peak responses after the first pulse. Positive and negative weights represented neurons with time-locked versus delayed first-pulse responses, respectively.

Mean ± SEM.

#### Supplementary tables

**Table S1** | Composition of complex particles. This table lists the compounds and their concentrations used in complex particles for larval zebrafish feeding assays.

**Table S2** | Detailed statistical results. Summary of statistical tests performed using Python.

**Table S3** | Statistical analysis decision guide. This table outlines the strategy for selecting appropriate statistical tests based on data type, distribution, and experimental design across the analyses in the study.

#### Supplementary videos

**Video S1** | Gut motility dynamics. Time-lapse video showing pixel intensity changes (Gaussian filtered) in the gut of a 9 dpf, unfed, wild-type zebrafish. See also Figure S2.

**Video S2** | Microgavage under a two-photon microscope. Infrared recording of a side gavage performed under a two-photon microscope, shown at 5X speed. The gavage needle travels ~1 mm from the mouth to the gut in ~5 minutes. Both the direct camera view and the prism view are used to position and guide the needle with precision, ensuring tissue integrity throughout the procedure.

**Video S3** | Encoding of gut stretch in vagal neurons. Two-photon calcium imaging of the left vagal ganglion in an 8 dpf zebrafish during repeated gut stretch. Neurons are color-coded by response: non-responsive (gray), suppressed (blue), and activated (red). The neutral 'water' stimulus was injected into the gut in five consecutive pulses (red vertical lines).

**Video S4** | Encoding of gut stretch in hindbrain neurons. Two-photon calcium imaging of the dorsal hindbrain in an 8 dpf zebrafish during repeated gut stretch. Neurons are color-coded by response: non-responsive (gray), suppressed (blue), and activated (red). The neutral 'water' stimulus was injected into the gut in five consecutive pulses (red vertical lines).

### Supplementary methods

#### *Generating transgenic zebrafish*

Tg(elavl3:H2B-GCaMP8m):

To generate the *Tg(elavl3:H2B-GCaMP8m)* line, we constructed a plasmid using a combination of PCR, restriction digestion, and NEBuilder HiFi DNA Assembly (E2621S, NEB). Three plasmids were used in the cloning strategy: (1) *Tol2-elavl3-TM-CaM-NES-TEV-N-AsLOV2-TEVseq-tTA*, (2) *Tol2-elavl3-H2B-GCaMP6s* (plasmid #59530, Addgene), and (3) *pGP-AAV-syn-FLEX-jGCaMP8m-WPRE* (plasmid #162378, Addgene). The *Tol2-elavl3-TM-CaM-NES-TEV-N-AsLOV2-TEVseq-tTA* plasmid was digested using NcoI-HF (R319S, NEB), NotI-HF (R3189S, NEB), and Quick CIP (M0525S, NEB) to remove the *TM-CaM-NES-TEV-N-AsLOV2-TEVseq-tTA* region. The H2B sequence was PCR-amplified from *Tol2-elavl3-H2B-GCaMP6s* using the following primers:

- Forward: ctagttctagagccaccatgCCAGAGCCAGCGAAGTCTG
- Reverse: ccatggtggcgaCTGGTGGATCCTTAGCGCTG

The jGCaMP8m sequence was PCR-amplified from *pGP-AAV-syn-FLEX-jGCaMP8m-WPRE* using:

- Forward: caccagtcgccaccatggctCATCATCACCATCATCACACGC
- Reverse: atgatctagagtcgcggccgcTCACTTCGCTGTCATCATTTGTACA

All three components (linearized backbone and PCR fragments) were ligated using NEBuilder HiFi DNA Assembly. The resulting plasmid was transformed into 5-alpha Competent *E. coli* (High Efficiency; C2987H, NEB) and purified using the Zyppy Plasmid Miniprep Kit (D4019, Zymo Research). Ligation and sequence integrity were verified by Sanger sequencing using primers near the ligation sites: tgttctgctggtagtggtag, tgattcatcagcagctgcg, and cgaaggacctagatcactatcatct.

Tg(neurod1:GCaMP8m-SV40):

To generate the *Tg(neurod1:GCaMP8m-SV40)* line, we used a two-step cloning approach. First, we received a *neurod1* promoter Gateway plasmid from the John Rawls lab (Duke University) and inserted it into the *pTol2-14xUAS-ChRmine-oScarlet-Kv2.1* backbone using Gibson assembly (NEBuilder HiFi DNA Assembly). The *neurod1* promoter was PCR-amplified using:

- Forward: ggcccccctcgagaagcttCCGGCATCAAACCGCCT
- Reverse: tgtgccatggtggcgaattcGTCGGAACTCTGCAAAGCGATAAAG

This produced a *neurod1:ChRmine-oScarlet-Kv2.1* intermediate plasmid. Next, we replaced the *ChRmine-oScarlet-Kv2.1* coding sequence with *GCaMP8m*, PCR-amplified from *pGP-AAV-syn-FLEX-jGCaMP8m-WPRE* using:

- Forward: CGACgaattcgccaccatggctcatcatcaccatcatcacacgc (adds Kozak sequence)
- Reverse: atgatctagagtcgcggccgctcacttcgctgtcatcatttgtaca

Assembly was again performed using NEBuilder HiFi DNA Assembly, and the resulting plasmid was transformed into 5-alpha Competent *E. coli* (High Efficiency), purified with the Zyppy Plasmid Miniprep Kit, sequence-verified, and prepared for injections.

Zebrafish microinjections:

At the 1-cell stage, zebrafish zygotes were injected with a DNA-Tol2 mRNA mixture. Injection mixtures, prepared in 0.5–1.5 mL tubes, contained:

- 1.5–3 µL of 25 ng/µL plasmid DNA (*elavl3:H2B-GCaMP8m* or *neurod1:GCaMP8m-SV40*)
- 1–2 µL of 100 ng/µL Tol2 mRNA
- 0.5–1 µL of phenol red (P0290, Sigma-Aldrich)

Tol2 mRNA was synthesized from the pCS2FA-transposase plasmid, linearized with NotI-HF, transcribed using the HiScribe SP6 RNA Synthesis Kit (E2070S, NEB), and purified using the RNA Clean & Concentrator-25 kit (R1017, Zymo Research). Each zygote was injected with ~1 nL of the mixture. F0 generations were screened for transgene expression via fluorescence.

##### *Preparation of complex particles for feeding experiments*

Liposomes were prepared using the method originally described by Barr and Helland (2007)[1] and modified by Hawkyard et al. (2015)[2]. To generate the liposome core solutions, water-soluble compounds of interest were dissolved in distilled water at a concentration of 75 mM in the pre-production mixture (referred to as the "mash"), resulting in an estimated final concentration of ~25 mM in the final complex particles (CPs) after production and washing. Sodium fluorescein (F6377, Sigma-Aldrich; 1 mg/mL in the core solution) was included in the liposome cores to enable fluorescence-based tracking of CP ingestion. Because fluorescein is water-soluble, it co-localizes with the encapsulated nutrients and serves as a proxy for their presence in the gut.

Core solutions were encapsulated in phospholipid membranes (Phospholipon 90 H, Lipoid) as described in Hawkyard et al. (2015)[2]. Full liposome formulations are provided in **Table S1**. These liposomes were then embedded into CPs as described in Hawkyard et al. (2019)[3]. Briefly, a 10% (w/v) sodium alginate (180947, Sigma-Aldrich) stock solution was prepared 24 hours in advance, and a 20% (w/v) cold-water fish gelatin (G7041, Sigma Aldrich) stock solution was prepared several hours before CP production by heating in a water bath (~45°C). The alginate and gelatin were added to the mash so that they comprised 10% and 33% of the total mash (by formula), respectively. The mixture was sonicated using an ultrasonic processor (FB505 equipped with a CL-334 wand, Fisherbrand) for one minute in four 15-second bursts. Liposome suspensions were then added to the mash at 57% (w/w) of the total mash (by formula), and the mixture was stirred for 1 minute using a stainless-steel spatula.

The final mash was loaded into a 60-mL syringe and sprayed into a 1% (w/v) $CaCl_2$ bath prepared in distilled water to induce gelation[3]. The resulting particles were filtered sequentially through 150-μm and 50-μm mesh screens to obtain CPs ranging from 50–150 μm in diameter.

##### *Gut-mediated feeding assay (additional details)*

At 8 dpf, the water in each petri dish was replaced in the morning to remove any residual paramecia. Zebrafish larvae were then transferred to new petri dishes for 24-hour fasting. For post-fixation washing, fish were transferred to 15 mL falcon tubes containing 0.1% Triton X-100 (9002-93-1, MP Biomedicals) in E3 medium and gently agitated on a shaker for 5 minutes per wash. This step was repeated three times to ensure thorough rinsing. A previously validated low-cost image acquisition method[4] using a smartphone and eyepiece adapter was employed for consistent image capture.

Images captured with the iPhone SE were transferred to a 2021 MacBook Pro (Apple) using the Photos app and exported as raw TIFF files. These were then imported into Fiji as individual files. Using the Freehand Selections tool, the gut of each fish was manually outlined. Selected gut areas were cropped without altering the image dimensions and saved as individual TIFF files. These cropped images were reimported into Fiji as a stack and converted to 8-bit grayscale. A conservative threshold (145–255) was applied to isolate fluorescent particles against a dark background. If this range failed to accurately capture visible signal or included unwanted areas, the threshold minimum was adjusted. Any deviations from the default threshold were recorded in a `notes.txt` file saved in the corresponding experimental folder. Thresholded regions were saved as binary masks and quantified using Fiji's Image Calculator function. Fluorescence intensity values were compiled into CSV files, with stimulus conditions manually labeled for each fish. These files served as input to a custom Python pipeline for batch analysis and visualization.

***Gut motility***

Larval zebrafish were raised following the same protocol as for the feeding behavior experiments (see "Gut-mediated feeding assay" in Methods). At 9 dpf, after a 24-hour fasting period, fish were split into fed and unfed groups. Fed fish were given paramecia for 45 minutes in the incubator, while unfed fish received no food. For imaging, fish were embedded on their sides in 1.5% (w/v) low-melting-point agarose (16520-050, Invitrogen). Gut motility was recorded under a 20X water-immersion two-photon objective (XLUMPLFLN20XW, Olympus) using infrared illumination from below (M780L3, Thorlabs) and captured with an infrared tissue camera (GS3-U3-41CNIR-C, Teledyne FLIR). The imaging field-of-view was centered on the boundary between the proximal and midgut, specifically focusing on the characteristic kink just posterior to the swim bladder. Recordings were acquired at 10 frames per second (fps) for 6 minutes using SpinView and saved as .avi files.

For analysis, the first 5 minutes of each video were imported into Fiji and converted to TIFF format. A mask was drawn to outline the gut and restrict the analysis to gut-specific movements. Total pixel changes between consecutive frames within the mask were calculated, and to account for variability in gut size across fish, these values were normalized to the total number of pixels within the mask, yielding a normalized pixel change trace. Since gut peristalsis occurs at low frequencies, traces were low-pass filtered using a $5^{th}$-order Butterworth filter with a 0.1 Hz cutoff frequency to eliminate high-frequency noise[5]. To mitigate the influence of random twitch events, the median of the filtered trace was used as the measure of average gut motility per fish. One fed $EEC^{DTA}$ fish was excluded from analysis as an outlier. This individual lacked observable twitching, displayed gross body deformation typical of dying fish, was flagged statistically using the ROUT method (Q=1%), and through visual inspection of the raw recording. The dataset containing this outlier is provided in the Prism file associated with Figure S2.

***Microgavage during two-photon imaging (additional details)***

Microgavage needle preparation (additional details):

Microgavage needles were prepared using a P-1000 pipette puller (Sutter Instrument), which was configured with the following settings: heat at ramp + 20, pull = 0, velocity = 90, and no time delay. We used a 2.5 mm square box filament, 4.5 mm wide (FB245B, Sutter Instrument), and performed a ramp test to optimize for this filament-glass combination. Pipettes were clipped immediately after pulling, and tip diameter was verified under a stereo microscope before continuing. To minimize tissue damage, needle tips were lightly fire-polished using a microforge (MF2, Narishige) to smooth sharp edges while preserving the opening.

Calibrating the gavage needle:

To calibrate the injection volume, 1–2 drops of mineral oil were placed onto a micrometer calibration slide. The needle was inserted into the oil, and the injector was set to 20 psi. The injection duration was adjusted to achieve consistent ejections of 2 nL per pulse. Due to slight variability in needle construction, such as the inner diameter at the bend, variation in tip diameter from fire polishing, and overall tip geometry, some needles exhibit capillary action that leads to unintended fluid suction. To test for this, the needle was left inserted in the injected stimulus droplet on the calibration slide for 2 minutes. If the droplet decreased in size, indicating capillary-driven reabsorption, the needle was discarded. ~10% of the fabricated needles passed this capillary action test and were used for experiments.

Microgavage during calcium imaging (additional details):

To ensure proper alignment during microgavage, the angle of the motorized micromanipulator (MP-865, Sutter Instrument) was adjusted so that the bent tip of the needle matched the angle at which the fish was embedded. Once the needle was inserted through the mouth, its left-right trajectory was adjusted every ~300 µm until reaching the esophagus, and then every ~100 µm until properly positioned in the proximal gut. Fish were monitored throughout for heartbeat stability and absence of struggling, which would indicate tissue injury.

Stimulus delivery was synchronized with the imaging software (PrairieView, Bruker) via voltage pulses. However, in 35.7% of experiments, a voltage signal failed to trigger the injector. These missed injections were non-systematic across trials and were identified by monitoring the injector's counter and by noting the absence of movement shifts in the imaging field of view (typically observed with each injection pulse). These discrepancies were recorded and accounted for during data analysis.

***Two-photon data processing and analysis (additional details)***

Parameters for vagal experiments:

For all vagal imaging experiments, calcium imaging data were processed using the following CaImAn parameters: `strides=100, overlaps=50, max_shifts=30, max_deviation_rigid=10, rf=25, stride=10, K=3, gSig=10, nb=2, merge_thr=0.999, rval_lowest=0.0, SNR_lowest=1.5, cnn_lowest=0.2, rval_thr=0.8, min_SNR=2.0,` and `min_cnn_thr=0.9`. To refine extracted components, we removed ROIs within 40 pixels of image borders and ROIs with raw peak fluorescence values below 80. When two ROIs had centers of mass within 20 pixels of each other, the one with the lower peak was excluded. Responses to injections were analyzed using a 15-frame (~11.5 s) baseline and a 13-frame (~10 s) post-injection window (**Figure S4B**).

For spatial registration across fish, we performed reverse alignment using SimpleITK[6–8]. Each fish's anatomical z-stack was first projected via summation and then aligned to the summed z-projection of a reference fish. The reference was chosen from a dataset with left vagal ganglion imaging. Therefore, anatomy stacks from right vagal ganglion recordings were flipped horizontally prior to alignment. To enhance contrast of anatomical landmarks, intensity values were normalized such that the minimum and maximum pixel intensities were set to 0 and 1000 for both reference and target images. Alignment was performed with `scalePenalty=150` and `iterations=(10000, 500)`. We manually defined the reference masks for vagal and posterior lateral line ganglia in Fiji. Given the relatively dim expression of *Tg(elavl3:H2B-GCaMP8m)* in vagal ganglia and the high inter-individual variability in ganglion morphology, spatial alignment could not reliably separate the posterior lateral line ganglion from the vagal ganglion. As a result, we took an inclusive approach, analyzing all neurons within the ganglionic region. Any neurons outside these regions were excluded from further analysis.

Parameters for hindbrain experiments:

For all hindbrain imaging experiments, calcium imaging data were processed using the following CaImAn parameters : `strides=80, overlaps=30, max_shifts=20, max_deviation_rigid=20, rf=25, stride=10, K=6, gSig=6, nb=2, merge_thr=0.999, rval_lowest=0.0, SNR_lowest=1.5, cnn_lowest=0.5, rval_thr=0.8, min_SNR=2.0,` and `min_cnn_thr=0.9`. To refine extracted components, we removed ROIs within 20 pixels of image borders and ROIs with raw peak fluorescence values below 40. When two ROIs had centers of mass within 10 pixels of each other, the one with the lower peak was excluded. Responses to injections were analyzed using a 15-frame (~11.5 s) baseline and a 20-frame (~15.3 s) post-injection window (**Figure S4B**).

For spatial registration, we selected the fourth imaging plane from each fish and computed a standard deviation projection across time. This projected image was aligned to the corresponding plane of a reference anatomical stack from a *Tg(elavl3:H2B-GCaMP8m)* fish. To enhance anatomical contrast, the minimum and maximum intensity were adjusted as follows: reference image = 0 to 1200 and target image = 400 to 1700. Alignment was performed with `scalePenalty=150` and `iterations=(10000, 500)`. To assign neurons to anatomical regions, we further aligned the GCaMP8m reference image to the Max Planck Zebrafish Brain ("mapzebrain") atlas[9] (https://mapzebrain.org). For this alignment, intensity values were adjusted to a minimum of 700 for the mapzebrain reference and 0 to 800 for the GCaMP8m reference. Alignment was performed with `scalePenalty=50` and `iterations=(10000, 500)`. Each neuron was then mapped to brain regions using mapzebrain region masks. Any neuron not assigned to a defined brain region was excluded from further analysis. Spatial plots in **Figures 4C** and **5C** show aligned neurons overlaid on the GCaMP8m reference image.

Temporal dynamic analyses:

For each neuron, both raw and denoised fluorescence traces were stored and normalized to a 0–1 range for consistent comparison (**Figure S4A**). To evaluate stimulus responsivity, we empirically determined the optimal pre-injection duration by calculating the standard error of the mean across baseline windows from 1 to 100 frames before each injection and identifying the elbow point (**Figure S4B**). A similar approach was used to determine the optimal post-injection window by plotting a histogram of peak times and identifying the elbow point (**Figure S4B**).

To assess stimulus-evoked responses, individual injection-aligned traces were extracted using the defined pre- and post-injection durations and normalized to the median of the pre-injection baseline. Each neuron's normalized traces were then averaged across all pulses. To classify neurons as "activated" or "suppressed", we compared the average post-injection trace to the baseline period. The standard deviation of the pre-injection baseline was calculated. A neuron was considered activated if the post-injection peak exceeded the baseline median by more than two standard deviations. Conversely, it was considered suppressed if the minimum post-injection value fell below the baseline median by more than two standard deviations. To prevent misclassification of non-biological artifacts with low variance, we applied an additional threshold, where activated neurons had to exceed a peak dF/F of 0.2, and suppressed neurons had to fall below -0.2. Next, we determined pulse-specific responsiveness by repeating this comparison for each injection individually. A neuron was classified as responsive if it responded to more than half of the total number of pulses (i.e., ≥3 out of 5).

For visualization, heatmaps of denoised traces were plotted and sorted by locating the maximum value of a sliding 10-frame window. For all other analyses, the raw traces were used. We quantified five key temporal response features (**Figure S4C**):

1. Peak global dF/F: Maximum post-injection value, normalized to the median baseline of the first injection (**Figure S4D**).
2. Peak local dF/F: Maximum post-injection value, normalized to the baseline of that specific injection (**Figure S4D**).
3. Baseline AUC: Total area under the curve (AUC) of the pre-injection baseline period, when the trace is normalized to the first injection baseline.
4. Time at half-maximum: Interpolated time before the peak when fluorescence reached 50% of the peak amplitude. If the peak occurred in the first frame post-injection, the value was set to 0 s. This metric was used instead of peak time to produce a more continuous distribution of response latencies.
5. Time to decay: Time from the peak until the signal returned to baseline. First, the peak frame was identified. Then, beginning from the next frame, a Wilcoxon signed-rank test compared the fluorescence value at each time point to the pre-injection distribution. If two consecutive non-significant frames occurred, the last significant frame was used to define the decay time. If the trace remained significant throughout the recording window (155 frames, ~119 s), that final frame was recorded. If the first two frames after the peak were non-significant, the decay time was assigned as one frame (~0.77 s).

Time at half-maximum and time to decay were both computed on traces normalized to their respective pre-injection baselines.

***Spatial distribution analyses of gut-responsive neurons***

Kernel density estimations (KDEs) were calculated from neuron coordinates aligned to the GCaMP8m reference (**Figures S6A, S8A, and S12A**). To evaluate topographical biases, we computed the AUC for

KDEs across defined spatial axes. Left vs. right and anterior vs. posterior were assessed by comparing AUCs from 0–250 μm vs. 250–500 μm along the corresponding axis. Dorsal vs. ventral differences were evaluated by comparing AUCs from 0–20 μm vs. 20–40 μm.

To quantitatively compare the spatial distributions of neurons between different classes, we calculated the Kullback-Leibler divergence (KLD), a measure of how one probability distribution diverges from another (**Figures S6B, S8B, and S12B**)[10]. For each pairwise comparison, neuron XYZ coordinates from the two groups of interest were used as input. We first generated continuous 3D probability distributions using Gaussian KDE for each population. To directly compare them, we created a shared 3D evaluation grid defined by the minimum and maximum XYZ coordinates across both distributions. This grid contained 100 equally spaced points per axis, yielding a total of $100^3$ = 1,000,000 evaluation points. KDE values were computed at each point and normalized to ensure both distributions summed to 1. The KLD was then computed as:

$$\mathrm{D}_{\mathrm{KL}}(\mathrm{P} \parallel \mathrm{Q}) = \sum_{\mathrm{i}} \mathrm{P(i)} \log \frac{\mathrm{P(i)}}{\mathrm{Q(i)}}$$

Since KLD is asymmetric (i.e., $D_{\mathrm{KL}}(P \parallel Q) \neq D_{\mathrm{KL}}(Q \parallel P)$), we computed it in both directions (i.e., $D_{\mathrm{KL}}(P \parallel Q)$ and $D_{\mathrm{KL}}(Q \parallel P)$), and averaged them to obtain a symmetrized KLD. When neuron counts were insufficient to produce stable KDEs, the analysis was omitted to avoid unreliable estimates.
